# Interictal Spikes in Alzheimer’s Disease: Preclinical Evidence for Dominance of the Dentate Gyrus and Cholinergic Control by Medial Septum

**DOI:** 10.1101/2023.04.24.537999

**Authors:** Christos Panagiotis Lisgaras, Helen E. Scharfman

**Affiliations:** Departments of Child & Adolescent Psychiatry, Neuroscience & Physiology, and Psychiatry, and the Neuroscience Institute, New York University Langone Health, 550 First Ave. New York, NY 10016; Center for Dementia Research, The Nathan Kline Institute for Psychiatric Research, New York State Office of Mental Health, 140 Old Orangeburg Road, Bldg. 35, Orangeburg, NY 10962

**Author notes:** **Corresponding author:** Christos Panagiotis Lisgaras The Nathan Kline Institute, Center for Dementia Research, 140 Old Orangeburg Rd. Bldg. 35, Orangeburg, NY 10962, Alternates.

**Keywords:** Alzheimer’s disease, amyloid precursor protein (APP), Presenilin 2, Down syndrome, Cholinergic neurons, Interictal spikes, Medial septum, Septohippocampal, Sleep

## Abstract

**HIGHLIGHTS:**

- Interictal spikes (IIS) occur in 3 mouse lines with Alzheimer’s disease features
- IIS in all 3 mouse lines were most frequent during rapid eye movement (REM) sleep
- The dentate gyrus showed larger IIS and earlier current sources vs. CA1 or cortex
- Chemogenetic silencing of medial septum (MS) cholinergic neurons reduced IIS during REM
- MS silencing did not change REM latency, duration, number of bouts or theta power

Interictal spikes (IIS) are a common type of abnormal electrical activity in Alzheimer’s disease (AD) and preclinical models. The brain regions where IIS are largest are not known but are important because such data would suggest sites that contribute to IIS generation. Because hippocampus and cortex exhibit altered excitability in AD models, we asked which areas dominate the activity during IIS along the cortical-CA1-dentate gyrus (DG) dorso-ventral axis. Because medial septal (MS) cholinergic neurons are overactive when IIS typically occur, we also tested the novel hypothesis that silencing the MS cholinergic neurons selectively would reduce IIS.

We used mice that simulate aspects of AD: Tg2576 mice, presenilin 2 (PS2) knockout mice and Ts65Dn mice. To selectively silence MS cholinergic neurons, Tg2576 mice were bred with choline-acetyltransferase (ChAT)-Cre mice and offspring were injected in the MS with AAV encoding inhibitory designer receptors exclusively activated by designer drugs (DREADDs). We recorded local field potentials along the cortical-CA1-DG axis using silicon probes during wakefulness, slow-wave sleep (SWS) and rapid eye movement (REM) sleep.

We detected IIS in all transgenic or knockout mice but not age-matched controls. IIS were detectable throughout the cortical-CA1-DG axis and occurred primarily during REM sleep. In all 3 mouse lines, IIS amplitudes were significantly greater in the DG granule cell layer vs. CA1 pyramidal layer or overlying cortex. Current source density analysis showed robust and early current sources in the DG, and additional sources in CA1 and the cortex also. Selective chemogenetic silencing of MS cholinergic neurons significantly reduced IIS rate during REM sleep without affecting the overall duration, number of REM bouts, latency to REM sleep, or theta power during REM. Notably, two control interventions showed no effects.

Consistent maximal amplitude and strong current sources of IIS in the DG suggest that the DG is remarkably active during IIS. In addition, selectively reducing MS cholinergic tone, at times when MS is hyperactive, could be a new strategy to reduce IIS in AD.

## INTRODUCTION

Abnormal electrical activity is gaining attention as an underappreciated contributor to Alzheimer’s disease (AD) pathophysiology (Palop and Mucke, 2009; Friedman et al., 2012; Scharfman, 2012; Edwards and Robertson, 2018; Lam and Noebels, 2020). Abnormal electrical activity has been reported in humans with AD (Lam et al., 2017; Vossel et al., 2017) and numerous mouse lines that simulate aspects of AD (Palop et al., 2007; Minkeviciene et al., 2009; Bezzina et al., 2015; Kam et al., 2016; Gureviciene et al., 2019; Beckman et al., 2020; Lisgaras and Scharfman, 2022a). Although outright seizures in AD may be rare (Hauser et al., 1986; Scarmeas et al., 2009; Friedman et al., 2012), intermittent abnormal electrical activity is much more frequent than seizures (Vossel et al., 2016), and may occur at earlier disease stages than seizures (Kam et al., 2016; Lisgaras and Scharfman, 2022a). Thus, targeting intermittent abnormal electrical activity in AD offers the opportunity to intervene early during AD progression when treatments are more likely to succeed (Sperling et al., 2011).

One of the earliest types of abnormal electrical activity in AD models are interictal spikes (IIS; (Palop and Mucke, 2009; Kam et al., 2016)) and high frequency oscillations (HFOs; (Lisgaras and Scharfman, 2022a). IIS in epilepsy are known to impair cognition (Kleen et al., 2010; Horak et al., 2017) and they are found to do so in AD as well (Sanchez et al., 2012; Vossel et al., 2016; Horvath et al., 2021). Indeed, anti-seizure medications (ASMs) can improve cognition in those AD patients that show IIS in the EEG (Vossel et al., 2021). Also, IIS in preclinical investigations of AD (Palop et al., 2007; Bezzina et al., 2015; Kam et al., 2016) and humans with AD (Vossel et al., 2016; Lam et al., 2017) are known to occur during sleep which suggests that they may impair sleep-dependent memory consolidation (Stickgold, 2005).

Despite the robust occurrence of IIS in AD, which areas of the brain may dominate the activity during IIS are currently unknown. One of the reasons for this gap in knowledge may be related to limitations in conducting studies that are invasive, using electrodes to penetrate different brain regions. In this context, minimally invasive approaches were instrumental in pointing to the temporal lobe as one of the areas where IIS and seizures can be robust (Vossel et al., 2013; Lam et al., 2017). A neuroimaging study in amnestic patients has provided even greater resolution on the possible sources of increased excitability within the hippocampus (Bakker et al., 2012). Indeed, the study (Bakker et al., 2012) showed that within the hippocampus, area CA3 and the dentate gyrus (DG) were the hotspots of increased excitability. Nevertheless, detailed electrophysiological recordings of IIS in AD are limited because EEG monitoring is not common clinical practice. In this context, preclinical investigations are valuable because the circuits involved in abnormal electrical activity could be examined in greater detail using invasive electrophysiological approaches.

We have shown that a new type of abnormal electrical activity in 3 mouse lines termed HFOs can be recorded primarily from the granule cell layer (GCL) of the DG (Lisgaras and Scharfman, 2022a). Here we ask whether the DG is one of the areas where IIS are most robust in amplitude compared to other regions such as area CA1 and overlying cortex. Thus, we analyzed IIS along the cortical-CA1-DG dorso-ventral axis. We recorded IIS in 3 different mouse lines each with different features relevant to AD. In all mouse lines, IIS were robust during rapid eye movement sleep (REM), a sleep stage when cholinergic tone is typically increased (Jasper and Tessier, 1971; Vazquez and Baghdoyan, 2001). We also found that IIS showed an amplitude gradient along the cortical-CA1-DG axis with IIS amplitude significantly greater in the DG GCL compared to hippocampal area CA1 and the overlying deep cortical layers. Notably, strong current sources were also prominent in the DG and typically followed by additional sources in area CA1 and overlying cortex. Thus, large IIS and early current sources in the DG suggest that the DG is remarkably active during IIS.

We next asked why IIS occur during sleep and specifically REM sleep. REM sleep is a sleep stage when cholinergic tone is significantly increased compared to non-REM (NREM) sleep (Jasper and Tessier, 1971; Vazquez and Baghdoyan, 2001; Pace-Schott and Hobson, 2002). In epilepsy, increased cholinergic tone is thought to facilitate epileptiform activity and seizures (Friedman et al., 2007; Maslarova et al., 2013; Mikroulis et al., 2018). In the Tg2576 mouse line, the cholinergic antagonist atropine reduced IIS in REM sleep, however, this effect was confounded by a parallel reduction in REM sleep duration (Kam et al., 2016). Thus, direct evidence for a possible contributing role of the cholinergic system to IIS generation is currently lacking.

Here we focused on selective targeting of the cholinergic neurons that give rise to the most dense projection to the hippocampus and are already implicated in memory (Hasselmo and Sarter, 2011; Zhang et al., 2021) and AD pathophysiology (Hedreen et al., 1984; Colom, 2006; Kim et al., 2019; Falangola et al., 2021), the medial septum (MS) cholinergic neurons. Importantly, we found that the rate of IIS during REM sleep was significantly reduced upon MS cholinergic silencing compared to baseline IIS rate during REM sleep. It is notable that the reduction in IIS rate during REM sleep was not due to reduced REM sleep duration, reduced number of REM bouts or reduced theta power compared to baseline REM sleep. Also, the latency to REM sleep was not significantly affected. Notably, two additional control experiments did not show any statistically significant effects on IIS rate.

In summary, our data show the remarkable dominance of the DG in IIS relative to CA1 and overlying cortex. The dominance of the DG was evident from the amplitude of the DG IIS and the robust and early sources in the DG by current source density analysis. The results are surprising given the DG is relatively quiet under normal conditions (Jung and McNaughton, 1993; Diamantaki et al., 2016; GoodSmith et al., 2017; Senzai and Buzsáki, 2017; Pofahl et al., 2021). We also present the novel finding that silencing MS cholinergic neurons selectively can significantly reduce IIS rates. This would be particularly relevant to disease stages when the MS cholinergic neurons appear to be hyperactive or at times of when there is increased cholinergic tone (DeKosky et al., 2002; Kam et al., 2016). The dominance of the DG and the cholinergic control of IIS are important because they could lead to new therapeutic strategies.

## MATERIALS AND METHODS

### I. Animals

A. Choice of mouse lines and terminology.

The mouse lines used in this study were selected based on their relevance to AD by exhibiting some characteristics found in AD. Below we refer to mice carrying a mutation (Tg2576^+/-^, Ts65Dn) as transgenic. We refer to Presenilin 2 (PS2) knockout (PS2KO^-/-^) mice as knockout. We refer to their wild type littermates (Tg2576^-/-^, PS2KO^+/+^, 2n, respectively) as controls.

The Tg2576 mouse line (stock #1349, Taconic Biosciences) was used because it is one of the most well-characterized mouse line of cerebral amyloid overexpression (Hsiao et al., 1996). It has features of AD such as progressive amyloid-β (Aβ) accumulation (Kawarabayashi et al., 2001; Ishii et al., 2014; Kam et al., 2016), cholinergic dysfunction (Apelt et al., 2002; Klingner et al., 2003; Ohno et al., 2004), sleep disturbances (Wisor et al., 2005; Kent et al., 2018), and cognitive impairment (Ohno et al., 2004; Jacobsen et al., 2006; Duffy et al., 2015; Kim et al., 2021). Tg2576 transgenic mice overexpress the human amyloid precursor protein (APP) using the hamster prion promoter and have the Swedish mutation (APP_Swe_) in APP isoform 695 (Hsiao et al., 1996). Tg2576 transgenic mice show memory deficits at the age of 3-4 months (Duffy et al., 2015) and Aβ plaques by 6-12 months of age (Hsiao et al., 1996; Kawarabayashi et al., 2001; Ishii et al., 2014; Kam et al., 2016). We used a total of 15 Tg2576 mice. Specifically, we used 10 Tg2576 transgenic mice (5 males, 5 females) and 5 Tg2576 control mice (3 males, 3 females). Tg2576 transgenic mice were ±1.9 months-old (range 1-19 months) at the time of recording. Tg2576 control mice were 5.3±3.1 months-old (range 1-18 months). No statistically significant age differences were found between control and transgenic Tg2576 mice (Mann-Whitney *U*-test, U=25, p>0.99). Although different ages were used, only 2 transgenic mice and 1 control were older than 14 months.

To test the role of MS cholinergic neurons in controlling IIS we crossed female homozygous choline acetyltransferase (ChAT) ChAT-Cre^+/+^ mice (Rossi et al., 2011) (stock #006410, The Jackson Laboratory) to male Tg2576 transgenic mice. This cross was used to allow the selective expression of viral vectors in MS cholinergic neurons as per previous studies (Vandecasteele et al., 2014; Desikan et al., 2018; Jin et al., 2019). A total of 7 ChAT-Cre::Tg2576 mice were used. Specifically, we used 4 males and 3 females. In addition, we used a total of 3 Tg2576 mice (1 male, 2 females) for the control experiments where CNO was injected in Tg2576 mice without AAV (Fig 9). We used 7 ChAT-Cre::Tg2576 transgenic mice and their ages were 6.6±2.3 months-old (range 2-16 months). The age of Tg2576 mice that were used as controls for the CNO-injected mice were ±1.1 months-old (range 3-7 months). There was no statistically significant difference between the ages of ChAT-Cre::Tg2576 transgenic mice and Tg2576 mice that were used as CNO controls (Mann-Whitney *U*-test, U=8.5, p=0.73).

Presenilin (PS2) knockout (PS2KO) mice were used to simulate reduced function of PS2, similar to mutations of the *PS2* gene in a form of familial AD with early onset (Lehmann et al., 2021). Mutations in *PS2* gene are linked to impaired Ca^2+^ homeostasis and increased neuronal excitability in mouse cell lines (Kipanyula et al., 2012) and cell lines derived from humans with familial AD (Zatti et al., 2004; Zatti et al., 2006; Galla et al., 2020). In addition, patients with mutation in the *PS2* gene experience seizures (Ezquerra et al., 2003; Jayadev et al., 2010b; Cortini et al., 2018) and cognitive impairment (Lleó et al., 2001; Ezquerra et al., 2003; Jayadev et al., 2010b; Zarea et al., 2016). Also, PS2KO mice show features of this form of AD such as hyperexcitability and behavioral impairments when aged and challenged in the kindling paradigm (Knox et al., 2023). PS2KO mice do not develop Aβ neuropathology (Herreman et al., 1999) which allowed us the valuable opportunity to study IIS independent of it. Notably, *PS2* gene encodes the predominant form of γ-secretase in microglia thereby allowing us to integrate non-neuronal contributors to AD hyperexcitability (Jayadev et al., 2010a). We used a total of 9 PS2KO mice. Specifically, we used 5 PS2KO (n=3 males, 2 females) and 4 controls (2 males, 2 females). PS2KO mice were 10.7±1.9 months-old (range 6-15 months) at the time of recording. PS2KO control mice were 10.7±1.9 months-old (range 6-15 months). No statistically significant age differences were found between control and knockout mice (unpaired t-test, t=0.009, df=6, p=0.99). Although different ages were used, only 2 knockout mice and 2 controls were older than 12 months.

The Ts65Dn mouse model of Down syndrome (stock #005252, The Jackson Laboratory) was used as an additional model that simulates features of AD because the majority of individuals with Down syndrome develop AD (Wisniewski et al., 1985; Head et al., 2012; Salehi et al., 2016). The overlap with AD includes progressive amyloid accumulation (Choi et al., 2009; Hamlett et al., 2016), cholinergic neurodegeneration (Cataldo et al., 2003; Salehi et al., 2006; Rozalem Aranha et al., 2023), and behavioral impairment (Holtzman et al., 1996; Escorihuela et al., 1998; Hamlett et al., 2016). Ts65Dn mice have a triplicated portion of the mouse chromosome 16, the ortholog of human chromosome triplicated in individuals with Down syndrome (Reinholdt et al., 2011). Thus, both mouse and human have triplication of the *APP* gene (Salehi et al., 2006). We used a total of 10 Ts65Dn mice. Specifically, we used 5 transgenic (2 males, 3 females) and 5 controls (3 males, 2 females). Ts65Dn transgenic mice were 13.2±2.9 months-old (range 2-24 months) at the time of recording and Ts65Dn control mice were 15.3±4.6 months-old (range 3-24 months). Although different ages were used, only 2 transgenic mice and 2 control were older than 22 months. No statistically significant differences were found between control and transgenic Ts65Dn mice (Mann-Whitney *U*-test, U=24.5, p=0.76). There were no statistically significant differences in age between the controls of different mouse lines (Kruskal-Wallis test, H(3)=3.89, p=0.14) and no differences in transgenic or knockout mice of the different mouse lines (Kruskal-Wallis test, H(3)=5.96, p=0.06).

#### A. B. Animal care

Tg2576 mice were bred on a C57BL6/SJL background (stock #100012, The Jackson Laboratory), PS2KO on a C57BL/6J background (stock #000664, The Jackson Laboratory) and Ts65Dn mice on a hybrid background (stock #003647, The Jackson Laboratory). Chat-Cre mice were maintained on a C57BL/6J background (stock #000664, The Jackson Laboratory).

All mice were fed Purina 5001 (W.F. Fisher) with water *ad libitum*. Cages were filled with corn cob bedding and there was a 12 hr light:dark cycle (7:00 a.m. lights on, 7:00 p.m. lights off). Genotyping was performed by the Mouse Genotyping Core Laboratory at New York University Langone Medical Center or in-house. All experimental procedures were performed in accordance with the NIH guidelines and approved by the Institutional Animal Care and Use Committee at the Nathan Kline Institute.

### II. Medial septum viral injections

Three weeks before the chemogenetic experiments, ChAT-Cre::Tg2576 mice were anesthetized with 3% isoflurane and then transferred to a stereotaxic apparatus where anesthesia was maintained with 1-2% isoflurane. Mice were placed on a heating pad (#50-7220F, Harvard Apparatus) with a rectal probe for control of body temperature. Next, buprenorphine (0.05 mg/kg) was injected subcutaneously (s.c.) to reduce discomfort. One burr hole was drilled above the MS (+0.7 mm Anterior-Posterior; A-P 0 mm Medio-Lateral; M-L) and 500 nL of adeno-associated virus (AAV) containing a modified designer receptor (hM4D) carrying an inhibitory signaling cascade (Gi; AAV5-hSyn-DIO-hM4D(Gi)-mCherry, Addgene) was stereotaxically injected over 10 min into the MS (−3.5 mm Dorso-Ventral; D-V, measured from the surface of the dura as before (Lisgaras and Scharfman, 2022b). The needle was then left in place for another 10 min and it was slowly retracted to avoid backflow of the injected virus. During the same surgery, one depth electrode was implanted in the left DG following procedures described below. Chemogenetic experiments were conducted 3 weeks after EEG surgery to allow for sufficient expression of the injected virus and recovery from EEG surgery.

### III. Electrode implantation

Mice were anesthetized, placed in a stereotaxic apparatus, and body temperature controlled as described above. Buprenorphine (0.05 mg/kg, s.c.) was injected as described above. Two burr holes were drilled above the cerebellum and subdural screw electrodes (2.54 mm length stainless steel, JI Morris) were placed and stabilized using dental cement (Lang Dental) to serve as a reference (−5.7 mm A-P, +1.25 mm M-L) and a ground (−5.7 mm A-P, −1.25 mm M-L). Next, a burr hole was drilled over the left hippocampus (−1.9 mm A-P, −1.2 mm M-L) and one 16-channel silicon probe (#A1×16, Neuronexus or #PLX-QP-3-16E-v2, Plexon) was implanted in the left dorsal DG (−1.9 mm A-P, −1.2 mm M-L, −1.9 mm D-V). The mice used for chemogenetic experiments were implanted with a single wire (instead of a silicone probe) in the left dorsal DG (−1.9 mm A-P, −1.2 mm M-L, −1.9 mm D-V). The single wire was 90 μm diameter stainless steel (California Fine Wire).

### IV. Wideband electrophysiological recordings

For local field potential (LFP) recording, mice were housed in a 21 cm x 19 cm square transparent plexiglass cage with access to food and water. A pre-amplifier (#C3335, Intan Techologies) was connected to the probe connector or the headset (in the case of mice recorded with a single wire) and then to a rotatory joint (Doric Lenses) via a lightweight cable to allow unrestricted movement of the mouse.

LFP signals were recorded at 2 kHz sampling rate using a bandpass filter (0.1-500 Hz) with either the Digital Lynx SX (Neuralynx) or an RHD interface board (#C3100, Intan Techologies) using the same settings. High frame rate video (>30 frames/sec) was recorded simultaneously using an infrared camera (#ac2000, Basler). Mice were continuously recorded for 3 consecutive days at a minimum.

### V. Chemogenetic silencing of MS cholinergic neurons and control experiments

For chemogenetic silencing of hM4D(Gi)-expressing MS cholinergic neurons and their axons, mice were injected intraperitoneally (i.p.) with CNO (3 mg/kg diluted in sterile saline; #BML-NS105-0005, Enzo Life Sciences) to activate the inhibitory Designer Receptors Exclusively Activated by Designer Drugs (iDREADDs). The CNO solution was made fresh before the start of each experiment. In the presence of CNO, hM4D(Gi) activates inward rectifying potassium channels that hyperpolarize hM4D(Gi)-expressing cells (Zhu et al., 2016). We used a 3 mg/kg dose of CNO because it is a standard dose used in the literature to inhibit diverse cell types with minimal to no off-target effects (Roth, 2016; Smith et al., 2016; Jendryka et al., 2019). Previous studies in mice have shown that blood plasma (Manvich et al., 2018) and brain (Jendryka et al., 2019) levels of CNO peak within 30 min after CNO injection, and the observed effects may last for hrs (Whissell et al., 2016).

We used 3 different groups to test the effect of MS chemogenetic silencing on IIS during REM sleep. In the experimental “CNO group,” we used ChAT-Cre::Tg2576 mice and injected CNO after a baseline. Effects were analyzed from 30-90 min after CNO injection. The waiting period is necessary for CNO levels to peak in the brain (Jendryka et al., 2019). A total of 4 ChAT-Cre::Tg2576 mice (2 males; 2 females) were used for these experiments. Notably, the mice were handled daily and were well-acclimated to the experimenter thus they did not show any signs of behavioral stress after injection that would likely influence the experiments.

We also used two control groups. For the “Saline group” we injected the same volume of sterile saline (0.9% NaCl) i.p. instead of injecting CNO. There were 3 ChAT-Cre::Tg2576 mice (2 males, 1 female). In an additional control group, we injected the same dose of CNO without AAV (“CNO no AAV group”). There were 3 Tg2576 transgenic mice (1 male, 2 females).

### VI. IIS detection, IIS amplitude analysis and Current source density analyses

#### A. IIS terminology, detection, and amplitude analysis

We use the term interictal spike (IIS) because we found previously that IIS just before a seizure in Tg2576 mice were similar in morphology to IIS in Tg2576 mice without seizures for several days (Kam et al., 2016). Notably, seizures in Tg2576 mice can be rare, requiring continuous video-EEG recordings >2 weeks for detection (Kam et al., 2016; Lisgaras and Scharfman, 2022a).

IIS in area CA1 were distinguished from sharp waves because sharp waves appeared well localized to CA1 stratum radiatum (SR; please see representative sharp wave in supplemental Fig 6 of Lisgaras and Scharfman, Epilepsia 2022 (Lisgaras and Scharfman, 2022a); see also (Buzsáki, 1986)), whereas IIS were present in every recording channel as shown in Figs 4-6. Also, IIS in the DG were distinguished from dentate spikes because dentate spikes are localized (Bragin et al., 1995; Buzsáki et al., 2003; Headley et al., 2017; Dvorak et al., 2021), and IIS were recorded in many brain areas. Also, dentate spikes are thought to be driven by the entorhinal cortex (Bragin et al., 1995; Dvorak et al., 2021) however, we cannot exclude the possibility that entorhinal projections contribute to IIS.

IIS were detected using the same criteria as described in our previous studies (Kam et al., 2016; Lisgaras and Scharfman, 2022a; Lisgaras et al., 2023). In brief, we included IIS that exceeded the mean of the baseline noise by more than 5 standard deviations (SDs). The baseline was taken from the same behavioral state as the state when IIS were recorded.

We also used a lower threshold (2 SDs) to make sure that IIS were not missed. This analysis was done in the cortex because the cell density is lower there than the CA1 or DG cell layers, making it more likely that small IIS might occur that might be missed if the threshold was too high. Notably, we did not find a statistically significant difference in cortical IIS rate when the threshold was 2 SDs vs. 5 SDs. Specifically, IIS rate for Tg2576 mice was 6.77±1.25 IIS/hr when 2 SDs was used as threshold vs. 6.25±1.30 IIS/hr when 5 SDs was applied as threshold (Wilcoxon signed rank test, p=0.25; n=4 mice). IIS rate in cortex of PS2KO mice was 0.09±0.03 IIS/hr when threshold was set at 2 SDs vs. 0.06±0.03 IIS/hr when threshold was 5 SDs (paired t-test, t*_crit_*=0.94, p=0.41; n=4 mice). Last, IIS rate in cortex of Ts65Dn mice using 2 SDs as threshold (0.07±0.03 IIS/hr) vs. 5 SDs (0.06±0.03 IIS/hr) was not significantly different (paired t-test, t*_crit_*=2.46, p=0.12; n=4 mice).

We analyzed IIS for a 24 hr recording period (except for the chemogenetics, as discussed below). During the 24 hr period, we noted which IIS occurred during wakefulness, slow-wave sleep (SWS), or rapid eye movement (REM) sleep. Awake and sleep stages were determined using the same criteria as in our previous studies and included simultaneous analyses of LFP and video records (Lisgaras and Scharfman, 2022a; Lisgaras and Scharfman, 2022b). In brief, awake state was defined by walking or other movement around the cage and the predominance of theta rhythm (4–12 Hz) with eyes open (when the mouse was facing the camera). SWS was characterized by the predominance of slow wave activity in the delta (<5 Hz) frequency range and eyes were closed with no movement of the body. REM sleep was defined by the predominance of theta rhythm (4–12 Hz) over delta and eyes closed with no movement of the body. Specifically, a theta/delta ratio of 2 was used as threshold to define REM sleep consistent with our previous studies (Lisgaras and Scharfman, 2022b; Lisgaras and Scharfman, 2022a). The rate of IIS per hr is reported separately for each behavioral state investigated (wakefulness, SWS, REM sleep). For spectral analyses, we used the time-frequency function in the RippleLab MATLAB application (Navarrete et al., 2016) and applied a 640-point window to visualize spectral power in the 0–15 Hz frequency range.

To quantify IIS amplitude across different recording channels (for Figs 4-6), the amplitude of IIS was measured at the time that IIS reached its maximum amplitude after crossing the detection threshold (defined above). IIS amplitude was measured for all detected IIS occurring within the 24 hr recording period. IIS amplitude measurements were done separately for each recording channel. We report the mean ± standard error of the mean (SEM) of IIS amplitude per recording channel as a mean per animal.

For MS chemogenetic experiments, IIS were detected using the same threshold as mentioned above (i.e., 5 SDs above the mean). Baseline IIS rate was calculated during two 1 hr-long recording periods preceding the CNO injection and the mean number of IIS per min is reported. As mentioned above, the 30 min after CNO injection was excluded from IIS analyses. After the 30 min waiting period following CNO injection, IIS were quantified from an 1 hr-long recording period. To determine if IIS levels returned to baseline, we analyzed IIS for an additional hr.

Theta (4–12 Hz) power during REM was quantified using Spike2 software (Cambridge Electronic Design, UK) using a Fast Fourier Transform (FFT) with a Hanning window. Latency to REM sleep was quantified by subtracting the time that the mouse entered the first REM sleep (as defined above) episode from the time that CNO was injected.

#### 1. B. Determination of electrode position

To determine the position of each electrode relative to the cell layers we used a combination of methods. While recording, we used electrophysiological landmarks as in our prior study.(Lisgaras and Scharfman, 2022a) In brief, the CA1 pyramidal layer was identified by the predominance of ripple oscillations in the 80-200 Hz frequency range and multi-unit activity (Oliva et al., 2016). SR was defined by the decrease in ripples and units, and the occurrence of sharp waves (Buzsáki, 1986). The DG was identified by the robust presence of fast activity in the gamma (∼80-150 Hz) frequency range (Fernández-Ruiz et al., 2021) and the GCL by the presence of multiunit activity.

In addition, histology was used to confirm that the electrode pierced the cortical surface, entered CA1 and ended in the DG GCL (see Fig 11A-B). Finally, we took advantage of the knowledge that silicon probes had electrodes that were 100 μm apart. Since probes were lowered so the first electrode was in the deep cortical layers (defined by the distance from the cortical surface), the other electrodes could be estimated as 100 μm intervals deep to the first electrode. The first electrode was placed in the deep cortical layers to allow simultaneous recordings in those layers, CA1 and the DG.

#### 1. C. Current source density analyses

Current source density analyses (CSD) were performed in NeuroScope2, an application in the MATLAB-based framework CellExplorer (Petersen et al., 2021). In order to quantify the maximal CSD intensity, CSD plots were converted in grayscale and imported in FIJI. Then the mean gray value was measured at the center of each current source in the DG, CA1 SR and cortex. The center of each source corresponded to maximal CSD intensity in colored CSD plots.

The latency to onset of each current source was determined after converting sources to white and the background to black. The conversion was done using a threshold that was based on current sources in cortex, CA1 and the DG. After establishing that cortical sources were the weakest (Fig 8B) compared to sources in CA1 and the DG, the next steps in setting the threshold used cortical sources. Thus, the weakest value within the cortical sources was defined (from Fig 8B), and then it was reduced by 2 SEM of the mean intensity of all cortical sources. This approach made it likely that even weak sources would be included. We then converted sources above threshold to white. After conversion, we found the following 3 main sources: DG, CA1 SR and cortex consistent with sources seen in colored CSD plots. Next, we measured the latency to each white source relative to a standard reference point (the onset of each CSD plot). The latency was measured using a free hand straight line tool in FIJI.

### VII. Histology

Mice were deeply anesthetized with isoflurane and then injected with an overdose of urethane (2.5 g/kg, i.p., Sigma Aldrich). Mice were transcardially perfused with 10 ml of room temperature (25 °C) saline, followed by 30 ml of cold (4 °C) 4% paraformaldehyde (Sigma Aldrich) dissolved in 0.1 M phosphate buffer (pH = 7.4). The brain was quickly removed and stored overnight in 4% paraformaldehyde. Coronal brain sections (50 μm) were cut using a vibratome (#Vibratome 3000, Ted Pella) and stored in cryoprotectant solution at 4 °C until use.

For Nissl stain, sections were mounted on 3% gelatin-coated slides and allowed to dry overnight. Then slides were dehydrated in increasing concentrations of ethanol (70%, 95%, 100%, 100%) for 2.5 min each, cleared in Xylene (Sigma Aldrich), and dehydrated again (100%, 100%, 95%, 70%) followed by hydration in double-distilled (dd) H_2_0 for 30 sec. Then sections were stained with 0.25% cresyl violet (Sigma Aldrich) in ddH_2_0 for 1.5 min followed by 30 sec in 4% acetic acid. Next, sections were dehydrated (70%, 95%, 100%, 100%), cleared in Xylene, and cover-slipped with Permount (Electron Microscopy Systems). Electrode placement was imaged using a microscope (#BX51, Olympus of America) equipped with a digital camera (#Infinity3-6URC, Olympus of America) at 2752×2192 pixel resolution and Infinity capture software (Olympus of America). The position of all electrodes was verified and reported in supplemental figures 2-4 of Lisgaras and Scharfman 2022.(Lisgaras and Scharfman, 2022a) An additional representative example of electrode targeting is shown in Fig 1B. To measure GCL thickness we traced the borders of the GCL using the freehand tool in FIJI as in our previous study (Lisgaras and Scharfman, 2022b). The border of the GCL and hilus and border of the GCL and inner molecular layer (IML) were defined by the point where adjacent cells became >1 GC soma width apart based on the traced borders of GCL as before (Lisgaras and Scharfman, 2022b). Analogous measurements were done for CA1 pyramidal cell layer (PCL) thickness. To measure thickness in both areas, we drew a straight line that was perpendicular to the length of the GCL or CA1 pyramidal layer as shown in Fig 7A-B.

**Figure 1:**
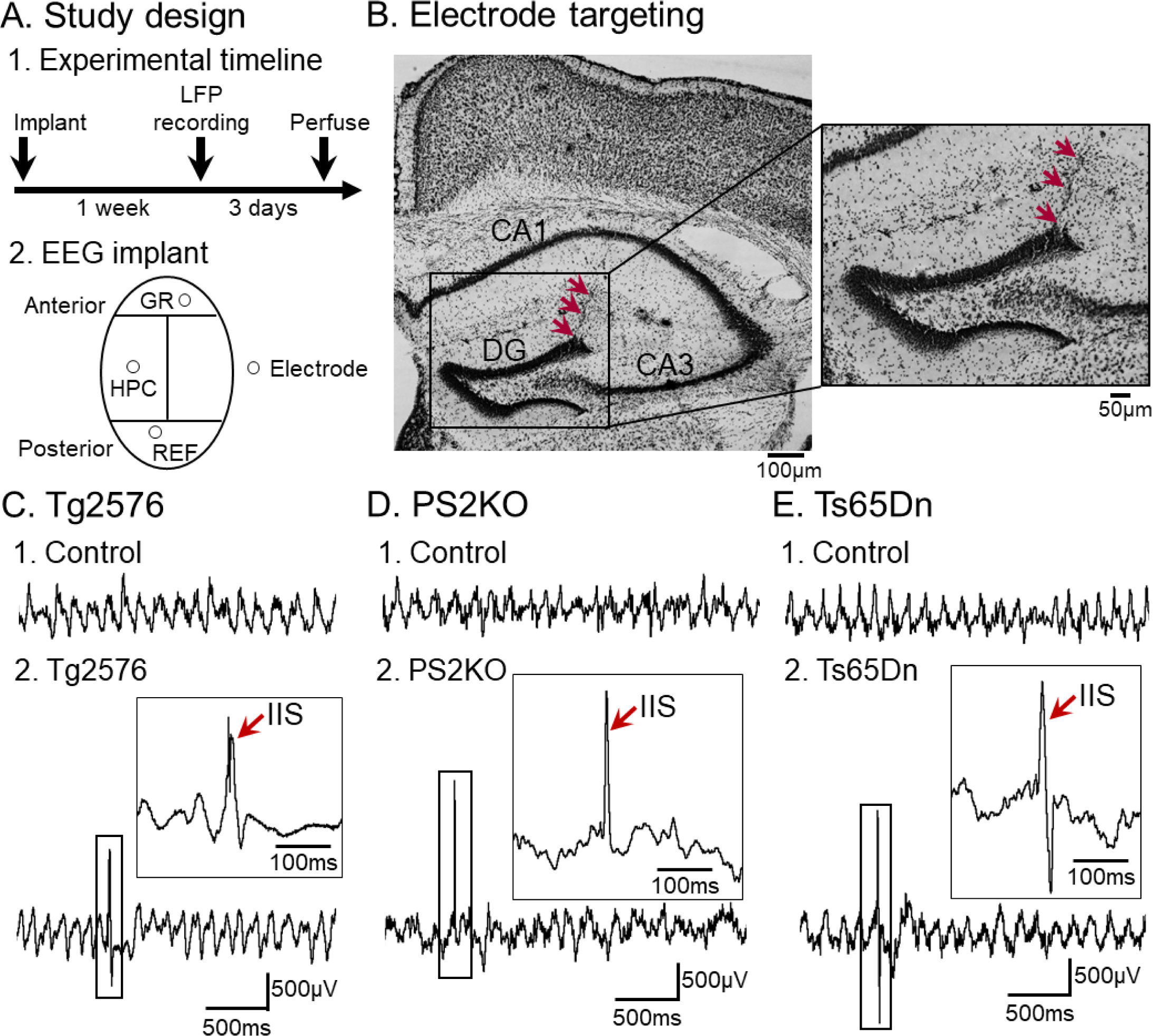
IIS in the DG is a robust EEG abnormality in three mouse lines. (A) 1. Experimental timeline of the study. Mice were implanted with a 16-channel silicon probe (for experiments without chemogenetics) or a depth electrode (for chemogenetics) and were allowed to recover from surgery for 1 week before the start of LFP and video recording. Video was also used as one component of the definition of behavioral state (see Methods). Following 1 week of recovery, LFP recording began and lasted for 3 consecutive days. After the end of the recording, animals were euthanized. 2. A diagram of the implant used for the EEG recordings. HPC=hippocampus; GR=ground; REF=reference. For simplicity, only one recording channel is shown for each mouse and the channel was in the GCL (note that IIS were detectable from additional areas – see Fig 3). (B) A representative Nissl-stained section confirming electrode targeting in the dentate gyrus (DG; red arrow). The electrode track is noted by red arrows. (C) Representative examples of recordings in the DG of a Tg2576 control and transgenic mouse. 1. Recordings in a control mouse. Note the absence of IIS. IIS were absent in all 5 control mice. 2. The same as 1 but the mouse was a Tg2576 transgenic. An IIS is shown (red arrow) and it is expanded in the inset. All 8 Tg2576 mice showed IIS. (D) Representative LFPs in the DG of a PS2KO control and knockout mouse. 1. Recordings in a control mouse. Note the absence of IIS. IIS were absent in all 4 control mice. 2. The same as 1 but the mouse was a PS2KO. An IIS is shown (red arrow) and it is expanded in the inset. All 5 PS2KO mice showed IIS. (E) Representative recordings in the DG of a 2n control and Ts65Dn transgenic mouse. 1. Recordings in the 2n control mouse. Note the absence of IIS. IIS were absent in all 5 control mice. 2. The same as 1 but the mouse was the Ts65Dn transgenic. An IIS is shown (red arrow) and it is expanded in the inset. All 5 Ts65Dn mice showed IIS.

To address AAV5-hSyn-DIO-hM4D(Gi)-mCherry viral expression in MS, 2 coronal brain sections adjacent to MS injection site (approximately +0.6 to +0.8 mm A-P, 0 M-L, −3.5 mm D-V) were mounted on 3% gelatin-coated slides and allowed to dry overnight under minimal light exposure. Then, the slides were coverslipped with Citifluor antifade mounting medium (Electron Microscopy Sciences), and micrographs were acquired at 4X using an epifluorescence microscope (#BX51, Olympus of America) with a digital camera (#Infinity3-6URC, Olympus of America) and Infinity capture software (Olympus of America). We visualized AAV5-hSyn-DIO-hM4D(Gi)-mCherry expression using mCherry fluorescence tag in the MS of ChAT-Cre::Tg2576 mice where Cre recombinase is known to be expressed in ChAT+ neurons (Vandecasteele et al., 2014; Desikan et al., 2018; Jin et al., 2019). Labeled cells were identified based on two criteria; their size was greater than 10 μm, consistent with a typical size of a neuron, and significantly greater mCherry signal intensity compared to the background. In the absence of a counterstain, we used other criteria to distinguish areas of interest (medial septum, MS; lateral septum, LS). First, we selected two sections corresponding to the location of these areas (approximately +0.6 to +0.8 mm A-P; −3.5 mm to −4.5 mm M-L, guided by a standard atlas (Franklin and Paxinos, 1997). Then the high background of MS was used to distinguish MS from LS (Fig 9E-F). We found robust AAV5-hSyn-DIO-hM4D(Gi)-mCherry expression in MS in all 4 mice we examined (typically 10-20 labeled cells per section). To address the specificity of AAV5-hSyn-DIO-hM4D(Gi)-mCherry expression for the MS, we compared the number of labeled cells per section (as well as mean per animal) in MS and LS. As shown in Fig 9G1-2, we found a significantly greater number of cells in MS compared to LS (typically 0-2 labeled cells per section) suggesting that viral expression was largely restricted to MS.

### VIII. Statistics

Data are presented as a mean ± SEM. Dots indicate individual data points. Statistical significance was set at p<0.05 and is denoted by asterisks on all graphs. Statistical comparisons that did not reach significance are designated by “ns” (not significant) in graphs and the exact p values are reported in text.

All statistical analyses were performed using Prism (Graphpad). To determine if data fit a normal distribution, the Shapiro-Wilk test was used. Comparisons of means for parametric data of two groups were conducted using unpaired two-tailed t-tests. Comparisons of data from one group before and after an experimental treatment were analyzed by paired two-tailed t-tests. When data did not fit a normal distribution, non-parametric statistics were used. The non-parametric test to compare two groups was the Mann-Whitney *U*-test. For comparisons of more than 2 groups, one-way ANOVA was used when data were parametric and Kruskal-Wallis for non-parametric data. When a statistically significant main effect was found by ANOVA, Bonferroni *post-hoc* tests were used with corrections for multiple comparisons and for Kruskal-Wallis, Dunn’s *post-hoc* tests were used. For ANOVA with repeated measures group comparisons with missing data points (Fig 5A), used a mixed-effects model and Holm-Šídák’s *post-hoc* tests. For parametric data with two main factors, two-way ANOVA was performed and for non-parametric data the Friedman test was used. When a statistically significant main effect was found, Tukey’s *post-hoc* tests were used with corrections for multiple comparisons. For correlation analyses we used linear regression. The Pearson correlation coefficient (*r*) was used for parametric data and Spearman *r* for non-parametric data.

## RESULTS

### I. IIS is a robust EEG abnormality in three mouse lines relevant to AD

To test whether IIS were robust in different mouse lines, mice were implanted with electrodes (Fig 1A1-2). The recordings used silicon probes for experiments without chemogenetics or depth electrodes for chemogenetic experiments. For GCL recordings, data were pooled from silicon probes and depth electrode recordings of the GCL. For other locations besides the GCL, data were from silicon probes only. A representative Nissl-stained section confirming electrode targeting in the DG is shown in Fig 1B.

After 1 week of recovery from surgery to implant electrodes, LFPs and video were recorded continuously for 3 consecutive days (Fig 1A1-2). We used 3 different mouse lines, each with a different causal mechanism and neuropathology (see Methods). For all mouse lines, age-matched controls (n=14) were recorded alongside the experimental (transgenic or knockout) mice (n=18). Representative LFPs from the DG of control and experimental mice are shown in Fig 1C-E.

We found that all transgenic and knockout mice showed IIS in the DG (n=8 Tg2576, n=5 PS2KO, n=3 Ts65Dn). An IIS recorded in a Tg2576 transgenic mouse is shown in Fig 1C2 and expanded in time in the inset. PS2KO and Ts65Dn transgenic mice also showed IIS and representative examples are shown in Fig 1D2 and Fig 1E2, respectively. Please note that IIS were also detectable in additional areas such as area CA1 and overlying cortex but were smaller (see Fig 3-6). We found that none of the age-matched controls showed IIS (n=5 Tg2576, n=4 PS2KO, n=5 Ts65Dn; Fig 1C1, 1D1, 1E1, respectively). Thus, IIS were an EEG event in experimental mice but not controls, and therefore were an abnormal electrical pattern.

### II. IIS are robust during REM sleep

We next asked when IIS occurred. We analyzed IIS during different behavioral states including wakefulness and sleep, which were distinguished based on established criteria (for details see Methods). In brief, wakefulness included times when mice were awake and immobile and times when they were actively exploring their home cage. Sleep was divided into SWS and REM using an elevated theta/delta ratio to define REM and no movement of the animal, as in our previous studies (Kam et al., 2016; Lisgaras and Scharfman, 2022a; Lisgaras and Scharfman, 2022b).

In all 18 transgenic and knockout mice (n=8 Tg2576, n=5 PS2KO, n=5 Ts65Dn) IIS occurred during REM sleep (Fig 2A-C, F) and were absent in wakefulness (Fig 2D) or SWS (Fig 2E). None of the 14 controls (n=5 Tg2576, n=4 PS2KO, n=5 2n) showed IIS in any behavioral state (Fig 2D-F).

**Figure 2:**
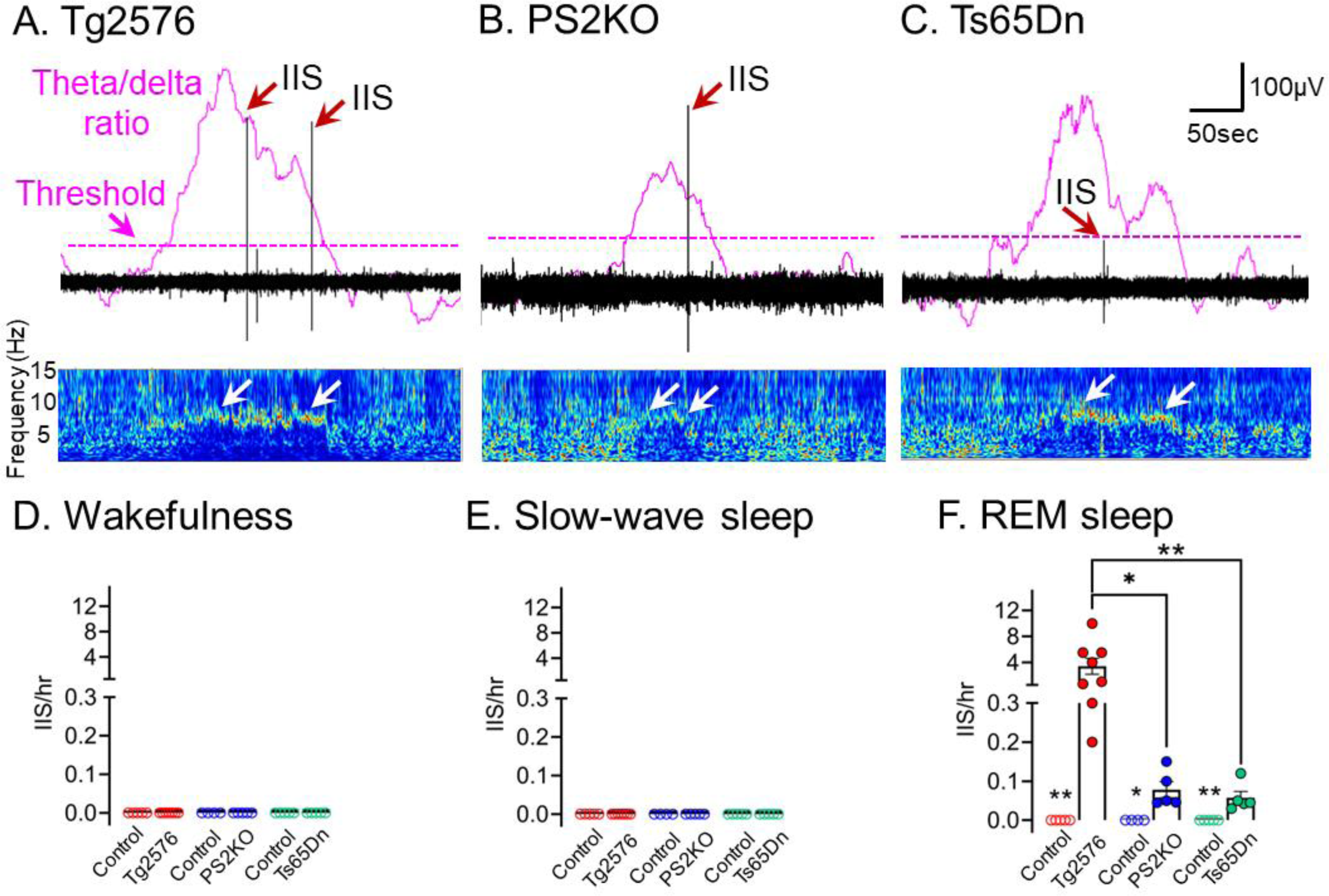
IIS rate in the DG is significantly increased during REM sleep. (A) Theta/delta ratio (purple line) and IIS (red arrows) in a Tg2576 transgenic mouse during sleep. The threshold for sleep of increased theta/delta ratio (2 times above the baseline) is denoted by a dotted purple line. Note that IIS occurred during periods of theta/delta ratio exceeding the threshold. Mice showed no movement during these periods determined by video (see Methods). Also note the increased theta power (white arrows) in the spectrogram during periods when IIS occurred, supporting the definition of this time as REM sleep. (B) Same as in A, but for a PS2KO mouse showing IIS during periods of increased theta/delta ratio. (C) Same as in B, but for a TS65Dn mouse. (D) IIS rate during wakefulness. None of the transgenic or control mice showed IIS during wakefulness. Red denotes Tg2576 control (n=5) and transgenic (n=8) mice. Blue denotes PS2KO control (n=4) and transgenic (n=5) mice, and green corresponds to 2n control (n=5) and Ts65Dn transgenic (n=5) mice. (E) Same as in B but for SWS sleep. None of the transgenic, knockout, or control mice showed IIS. (F) Same as in C but IIS were analyzed during REM sleep. IIS in Tg2576 transgenic mice were most frequent compared to PS2KO (p=0.01) or Ts65Dn (p=0.005) transgenic mice. Note that none of the controls showed IIS (n=14). Also, IIS were significantly more frequent in Tg2576 transgenic vs. control mice (Mann-Whitney *U*-test, U=0, p=0.001), PS2KO vs. control mice (Mann-Whitney *U*-test, U=0, p=0.01) and Ts65Dn transgenic vs. control mice (Mann-Whitney *U*-test, U=0, p=0.007). The mean number of IIS detected during a 24hr session was: Tg2576: 81.54±29.70 events (range: 5-240); PS2KO: 1.87±0.05 events (range: 1-4); Ts65Dn: 1.38±0.04 events (range: 1-3).

We next compared IIS rate between the different mouse lines. A Kruskal-Wallis test revealed that IIS rate was significantly different between mouse lines during REM sleep (Kruskal-Wallis test, H(3)=12.79, p<0.0001; Fig 2F). *Post-hoc* comparisons confirmed that IIS rate was significantly higher in Tg2576 transgenic mice vs. PS2KO (p=0.02; Fig 2D) or Ts65Dn (p=0.002; Fig 2D) transgenic mice.

Notably, we found no sex differences in the IIS rate of Tg2576 mice (unpaired t-test, t*_crit_*=0.53, p=0.61; 4 males, 4 females). Since we did not find a statistically significant interaction between mouse line and sex using two-way ANOVA (F(2, 10) = 0.11, p=0.89), and sample sizes of males and females in other mouse lines were small, we pooled males and females from all mouse lines for additional analyses (9 males, 9 females). Again, we found no statistically significant differences between males and females in IIS rate (Mann-Whitney *U*-test, U=31.5, p=0.45).

### III. Multi-area recording of IIS rate along the dorso-ventral cortical-CA1-DG axis

Our electrodes recorded LFPs along the dorso-ventral cortical-CA1-DG axis, allowing us to quantify IIS rate and amplitude (see Figs 4-6) in areas besides the DG (Fig 3A). To quantify IIS rate in cortex and area CA1 we selected the largest IIS in cortex, CA1, and the DG which corresponded to the deep layers, CA1 cell layer, and DG cell layer, respectively. The position of the electrodes was estimated based on electrophysiological landmarks, spacing between electrodes and post-mortem histological reconstruction of electrode targeting (for details see Methods).

**Figure 3:**
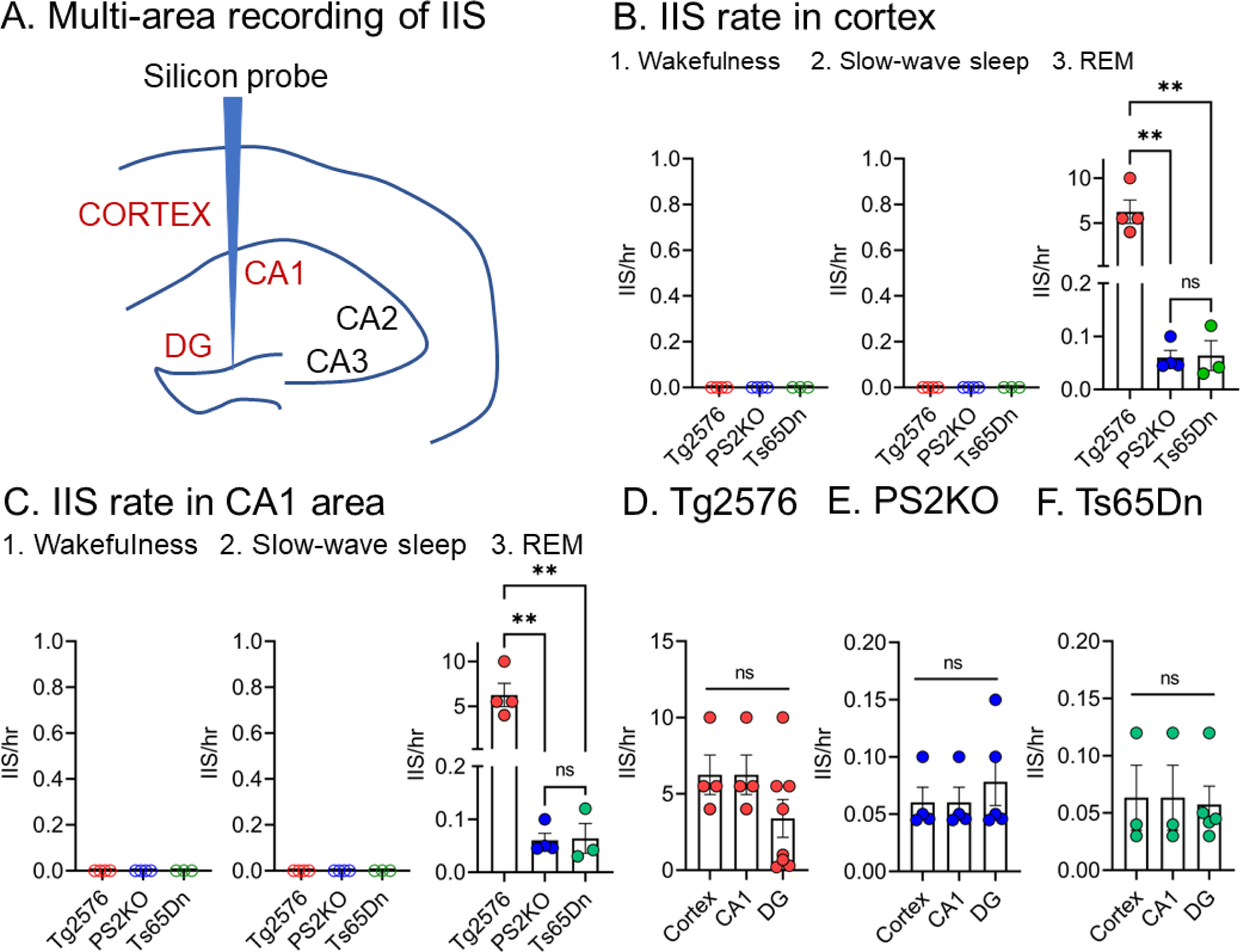
IIS rate in cortex, CA1 and the DG. (A) Schematic of electrode placement used for multi-area recording of IIS. IIS were recorded simultaneously in cortex, area CA1 and the DG using 16-channel silicon probes (n=4 Tg2576, n=4 PS2KO, n=3 Ts65Dn). In an additional cohort of mice, IIS were recorded using a depth electrode targeting the DG (n=3 Tg2576, n=2 Ts65Dn). (B) IIS rate per hr in cortex of Tg2576 (red; n=4), PS2KO (blue; n=4) and Ts65Dn (green; n=3) mice. 1. The IIS rate is shown for wakefulness. 2. The same as in 1 but for SWS. 3. The same as in 1, 2 but for REM sleep. One-way ANOVA revealed that IIS rate was significantly different between mouse lines during REM sleep (F(2, 8)=19.25, p=0.0009). *Post-hoc* comparisons showed that IIS rate was significantly higher in Tg2576 transgenic mice vs. PS2KO (p=0.002) or Ts65Dn (p=0.002) mice, but not between PS2KO and Ts65Dn mice (p=0.99). ns = not significant. (C) Same as in B but IIS rate was measured in CA1. Similar to other areas, IIS rate in CA1 was also robust in REM. There was a statistically significant difference in IIS rate depending on the mouse line (one-way ANOVA; F(2, 8)=19.25, p=0.0009). *Post-hoc* comparisons showed a statistically greater IIS rate in Tg2576 vs. PS2KO mice (p=0.002), Tg2576 vs. Ts65Dn (p=0.002) but not between PS2KO and Ts65Dn (p=0.99). (D) Multi-area comparison of IIS rate in Tg2576 transgenic mice. There was no statistically significant difference in IIS rate between the 3 different regions (one-way ANOVA; F(2, 13)=1.68, p=0.22). (E) Same as in D but for IIS rate in PS2KO mice. Similarly to Tg2576, IIS rate in PS2KO mice was not significantly different between the 3 areas (Kruskal-Wallis test, H(3)=0.35, p=0.35). (F) Same as in E but for IIS rate in Ts65Dn mice. IIS rate between the 3 areas was not significantly different (Kruskal-Wallis test, H(3)=0.86, p=0.31).

Similar to the results in the DG (Fig 2), IIS in cortex were not detected during wakefulness (Fig 3B1) or SWS (Fig 3B2) but were robust in REM (Fig 3B3). One-way ANOVA revealed that IIS rate was significantly different between mouse lines during REM sleep (F(2, 8)=19.25, p=0.0009; Fig 3B3). *Post-hoc* comparisons showed that IIS rate was significantly higher in Tg2576 transgenic mice vs. PS2KO (p=0.002; Fig 3B3) or Ts65Dn (p=0.002; Fig 3B3) mice, but not between PS2KO and Ts65Dn mice (p=0.99). IIS rate in area CA1 was also robust in REM (Fig 3C3). One-way ANOVA showed that the CA1 IIS rate was significantly different between mouse lines (F(2, 8)=19.25, p=0.0009; Fig 3C3). Like cortex, *post-hoc* comparisons revealed that CA1 IIS rate was significantly greater in Tg2576 mice compared to PS2KO (p=0.002), or Ts65Dn (p=0.002) but PS2KO and Ts65Dn mice were not different (p=0.99).

Last, we compared IIS rate between the 3 different areas (cortex, area CA1 and the DG) to determine if IIS rate was different based on the area that IIS were recorded from. Notably, we found that IIS rate was similar across the different areas in Tg2576 mice (one-way ANOVA, F(2, 13)=1.68, p=0.22; Fig 3D), PS2KO mice (Kruskal-Wallis test, H(3)=0.35, p=0.35; Fig 3E) and Ts65Dn mice (Kruskal-Wallis test, H(3)=0.86, p=0.31; Fig 3F). The results are consistent with the fact that IIS in mice simulating AD are recorded throughout the brain when they occur (Palop et al., 2007; Minkeviciene et al., 2009; Bezzina et al., 2015; Kam et al., 2016; Gureviciene et al., 2019; Beckman et al., 2020; Lisgaras and Scharfman, 2022a).

### IV. IIS show an amplitude gradient along the cortical-CA1-DG axis and maximal amplitude within the DG

To test whether there was a location along the cortical-CA1-DG axis where IIS were consistently maximal, transgenic and knockout mice were implanted with a 16-channel linear silicon probe (see Methods) with the tip ending in the dorsal DG. The silicon probe spanned the deep layers of the cortex to the DG GCL to capture the maximal area.

A representative example of simultaneous recordings during an IIS from a Tg2576 transgenic mouse is shown in Fig 4A. We found that IIS were detectable throughout the cortical-CA1-DG axis. Insets show IIS recorded in the GCL, CA1 PCL or a deep cortical layer. One-way ANOVA revealed a statistically significant difference in IIS amplitudes for the different recording channels (F(14, 30)=2.43, p=0.02, n=4 mice; Fig 4B). *Post-hoc* comparisons confirmed that IIS amplitude was significantly larger in the GCL vs. CA1 SR (p=0.003; Fig 4B), CA1 PCL (p=0.0003; Fig 4B) or deep cortical layers (p=0.0004; Fig 4B).

**Figure 4:**
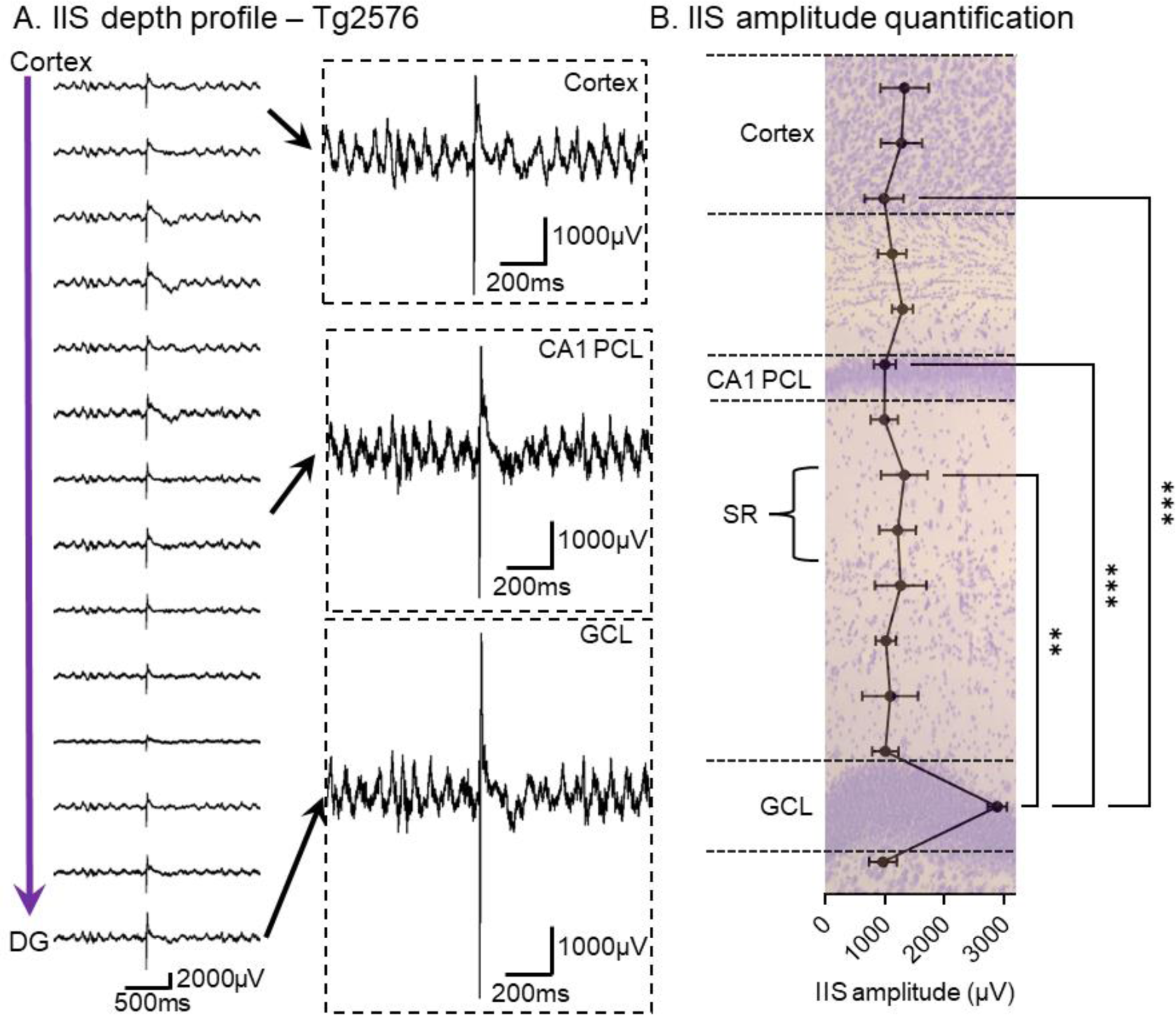
IIS amplitude is maximal in the GCL of Tg2576 transgenic mice. (A) A representative IIS recorded from a Tg2576 transgenic mouse. The recording channels are shown with the most superficial (closer to cortex) at the top and deeper channels (closer to the DG) at the bottom. Note that the IIS was detectable from all recording channels. Insets show an IIS in a deep cortical layer (top), CA1 PCL (center) and the GCL (bottom). In Figs 4-6, please note that 16-channel probes were used, but recordings from <16 channels are shown because some channels were noisy or not functional. (B) Quantification of IIS amplitude along the cortical-CA1-DG axis of Tg2576 transgenic mice (a total of 156 IIS were analyzed in 4 mice during 24 hr). The different layers were determined based on several criteria (see Methods). One-way ANOVA revealed a statistically significant difference in IIS amplitudes for different recording channels (F(14, 30)=2.43, p=0.02). *Post-hoc* comparisons confirmed that IIS amplitude was significantly larger in the GCL vs. CA1 SR (p=0.003), CA1 PCL (p=0.0003), or deep cortical layers (p=0.0004). In Figs 4-6, the brain section shown in the background was taken from a normal animal and it is used only for illustration.

We next asked whether the maximal IIS amplitude in the GCL of Tg2576 transgenic mice was generalizable to other mice that simulate aspects of AD. To that end, we implanted PS2KO mice and we found that IIS also showed a statistically significant amplitude gradient along the cortical-CA1-DG axis (Mixed-effects model, F(1.67, 4.88)=68.30, p=0.0003, n=4 mice; Fig 5A). *Post-hoc* comparisons confirmed that IIS amplitude was significantly larger in the GCL compared to CA1 pyramidal layer (p=0.009; Fig 5B) or deep cortical layers (p=0.009; Fig 5B). IIS in PS2KO mice were recorded in REM sleep, like Tg2576 mice.

**Figure 5:**
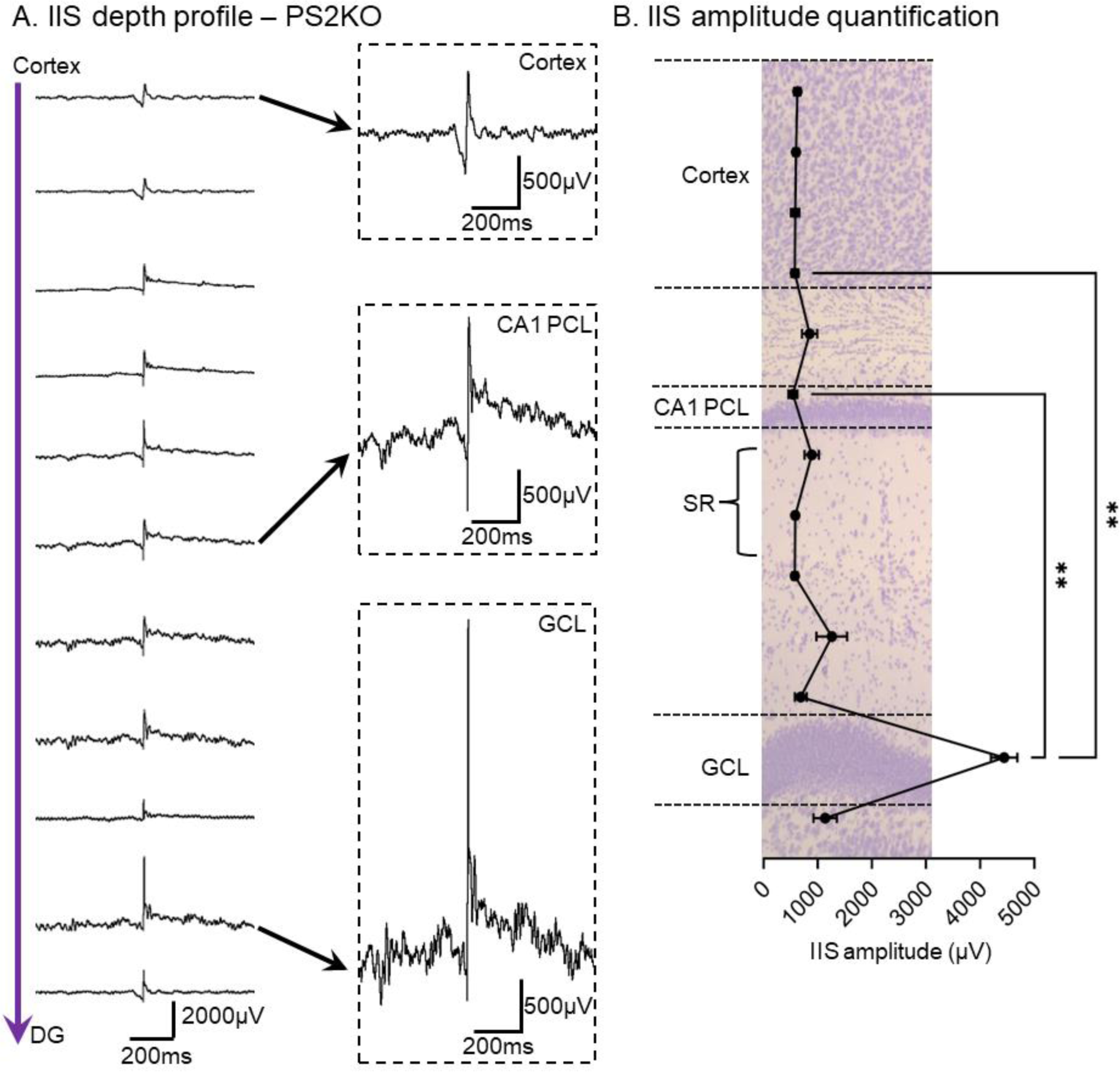
IIS amplitude is maximal in the DG of PS2KO mice. (A) A representative IIS recorded from a PS2KO mouse is shown. Similar to Tg2576 transgenic mice, IIS in PS2KO mice were detectable from all recording channels. Insets show an IIS in a deep cortical layer (top), CA1 pyramidal layer (center) and the GCL (bottom). (B) Quantification of IIS amplitude along the cortical-CA1-DG axis of PS2KO mice (a total of 22 IIS were analyzed in 4 mice during 24 hr). One-way ANOVA analysis revealed a statistically significant difference in IIS amplitude and recording channel (Mixed-effects model, F(1.67, 4.88)=68.30, p=0.0003). *Post-hoc* comparisons confirmed that IIS amplitude was significantly larger in the GCL vs. CA1 PCL (p=0.009) or deep cortical layers (p=0.009).

In a third mouse line, the Ts65Dn mouse, we also found a statistically significant IIS amplitude gradient along the cortical-CA1-DG axis (Friedman test (12)=25.85, p=0.007, n=3 mice; Fig 6A-B). *Post-hoc* comparisons revealed a significant increase in IIS amplitude in the GCL vs. CA1 PCL (p=0.04; Fig 6B) or deep cortical layers (p=0.03; Fig 6B).

**Figure 6:**
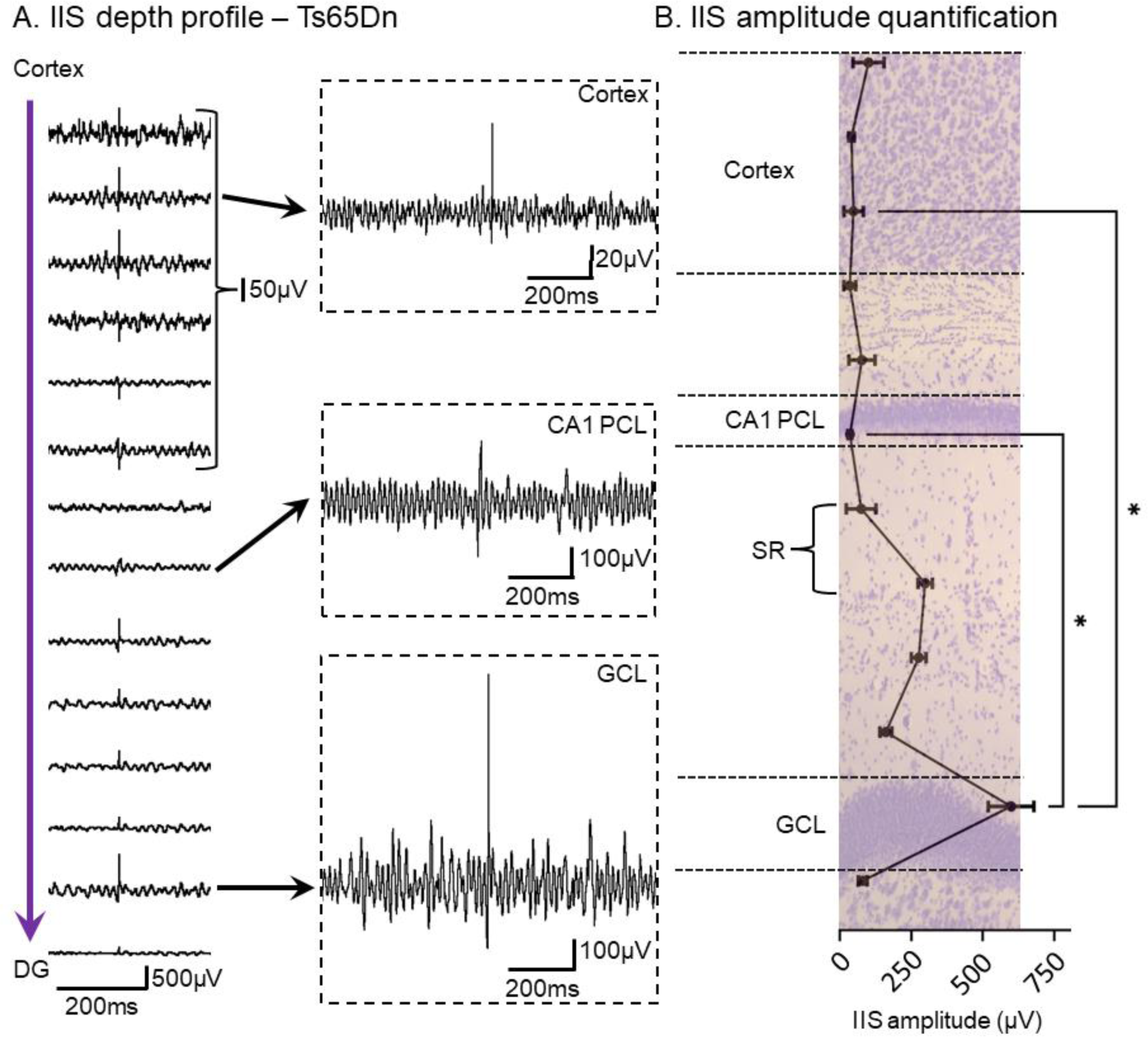
IIS amplitude is maximal in the DG of Ts65Dn transgenic mice. (A) A representative IIS recorded in a Ts65Dn mouse is shown. Like Tg2576 and PS2KO mice, IIS in Ts65Dn mice were detectable from all recording channels. Insets show IIS in a deep cortical layer (top), CA1 PCL (center) and the GCL (bottom). (B) Quantification of IIS amplitude along the cortical-CA1-DG axis of Ts65Dn transgenic mice (a total of 16 IIS were analyzed in 3 mice during 24 hr). A Friedman test revealed a statistically significant difference for IIS amplitude and recording channel (Friedman test (12)=25.85, p=0.007). Post-hoc comparisons confirmed that IIS amplitude was significantly larger in the GCL vs. CA1 PCL (p=0.04) or deep cortical layers (p=0.03).

### V. IIS amplitude and cell layer thickness are not linearly correlated

It was notable that the largest IIS were detected near the GCL. We next asked whether IIS amplitude was correlated to areas where cell density is typically increased i.e., GCL vs. CA1 PCL. The rationale was that a dense cell layer would be more likely to exhibit large LFPs, however it is possible that cell density may not be the only contributing factor controlling IIS amplitude. One reason is that large IIS could be recorded in areas where cells were not dense (Figs. 4-6), although this could be related to volume conduction. To quantitatively address this issue, we compared IIS amplitude with cell layer thickness (Fig 7A-B). Here cell layer thickness is used as a proxy of cell density although it is recognized it is not the same. Nevertheless, cell layer thickness is probably a good proxy for cell density, because there is a fairly homogeneous density of cells in the GCL and CA1 PCL.

**Figure 7:**
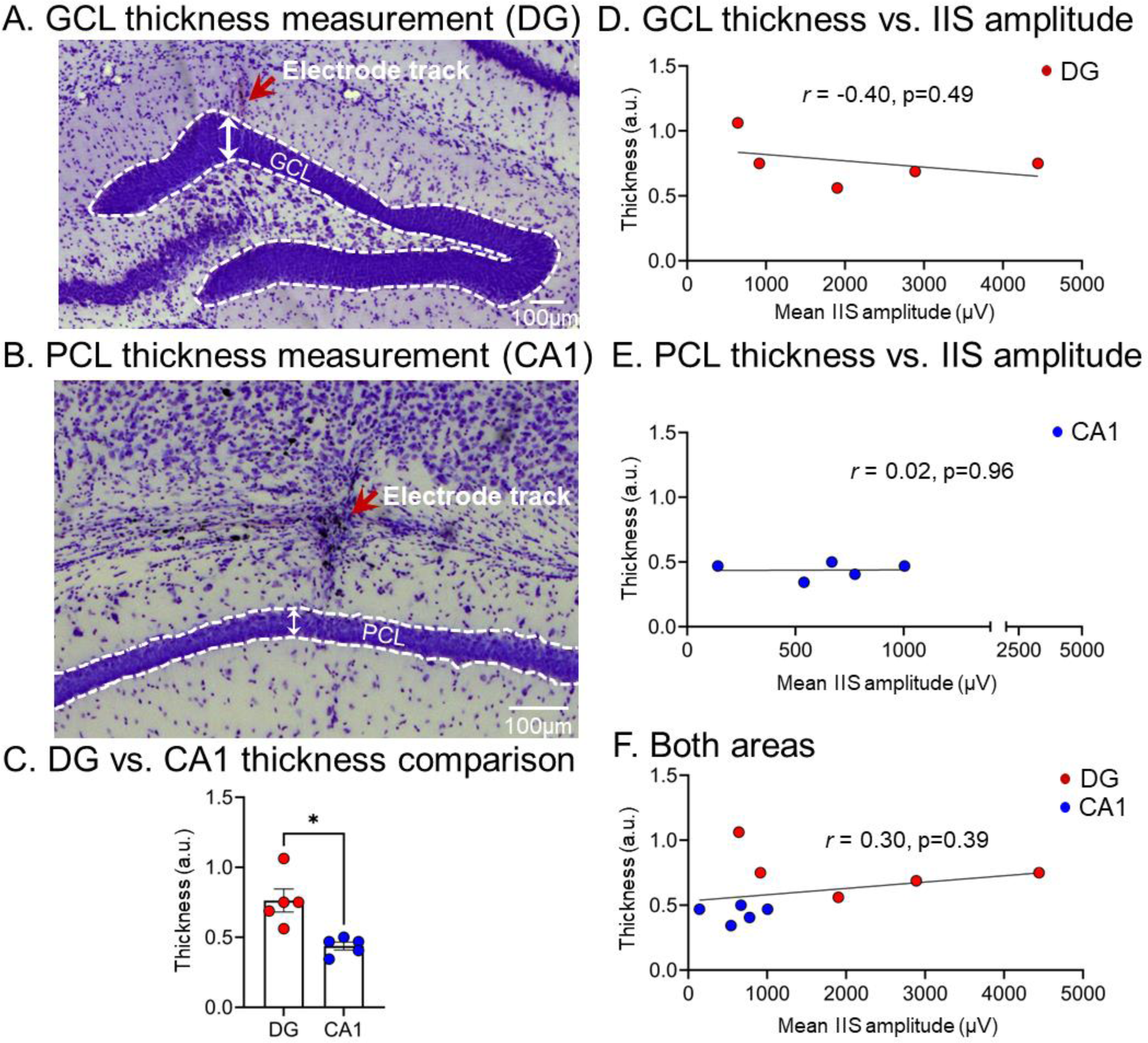
IIS amplitude and principal cell layer thickness are not linearly correlated. (A) Quantification of GCL thickness in the DG. For GCL thickness (width in arbitrary units, a.u.), we used the straight-line tool in FIJI to measure the thickest part of the upper blade. The GCL/hilar border and GCL/IML border was defined as before (Lisgaras and Scharfman, 2022b; see also Methods). (B) Same as in A, but for CA1 PCL thickness. The borders were defined in an analogous manner. (C) Comparison between GCL and PCL thickness. GCL thickness was significantly greater than PCL thickness (paired t-test, t*_crit_*=4.13, p=0.01; n=5 mice). (D) There was no statistically significant linear correlation between GCL thickness and mean IIS amplitude recorded in the GCL (Pearson *r* = −0.41, p=0.49). (E) There was no statistically significant linear correlation between the CA1 PCL thickness and mean IIS amplitude recorded in the CA1 PCL (Pearson *r* = 0.02, p=0.96). (F) Measurements from both the GCL and PCL were pooled. There was no statistically significant linear correlation (Pearson *r* = 0.30, p=0.39) between cell layer thickness and mean IIS amplitude.

Mean IIS amplitude and thickness in the GCL and CA1 PCL showed that the GCL was significantly thicker compared to the CA1 PCL, as expected (paired t-test, t*_crit_*=4.13, p=0.01; n=5 mice, 2 Tg2576, 2 PS2KO, 1 Ts65Dn; Fig 7C). There was no correlation between either the GCL thickness and mean IIS amplitude in the GCL (Pearson *r* = −0.41, p=0.49; Fig 7D) or CA1 PCL thickness and mean IIS amplitude in the PCL (Pearson *r* = 0.02, p=0.96; Fig 7E). When measurements of cell layer thickness from the DG and CA1 were pooled, again we did not find a statistically significant correlation of cell layer thickness and IIS amplitude (Pearson *r* = 0.30, p=0.39; Fig 7F). Thus, it seems unlikely that cell layer thickness alone, and probably cell density, accounts for the large IIS amplitude we recorded in the GCL. In summary, the large amplitude of the GCL LFP could be related to the cell density there, and this is discussed in the Limitations section of the Discussion. However, there are arguments against this idea, and moreover, data from CSDs support a dominance of the DG in other ways relative to overlying CA1 and cortex (see below).

### VI. IIS show early and strong current sources in the DG and additional sources in CA1 and cortex

We next used current source density (CSD) analysis to map the distribution of sinks and sources of IIS. In all mice we tested (n=6 mice: 2 Tg2576; 2 PS2KO; 2 Ts65Dn) we found that the DG always showed a current source during IIS (Fig 8A1-4). Notably, current sources in the DG remained robust over different days in the same mouse (compare Fig 8A1 vs. 8A2). However, additional current sources were found in CA1 SR (black arrows; Fig 8A1-4) and cortex (white arrows; Fig 8A1-4). To determine whether one of these sources showed stronger CSD intensity than others, we measured the maximal intensity of current sources in the DG, CA1 SR and cortex (Fig 8B). We found a statistically significant difference in maximal CSD intensity between the different areas (Friedman test (3)=8.09, p=0.009; Fig 8B). *Post-hoc* comparisons showed the DG source was stronger than the cortical source (p=0.01; Fig 8B) and there was a trend for the DG source to be greater than the CA1 SR source (p=0.06; Fig 8B). When comparisons were made using all IIS instead of the mean per animal, all comparisons were statistically significant (Friedman test (3)=27.08, p<0.0001). Thus, *post-hoc* comparisons showed a significantly greater CSD intensity in the DG compared to cortex (p<0.0001) and the DG was greater than CA1 (p=0.0001). These data suggest that the DG current sources are strongest and support the idea that the DG is dominant.

**Figure 8:**
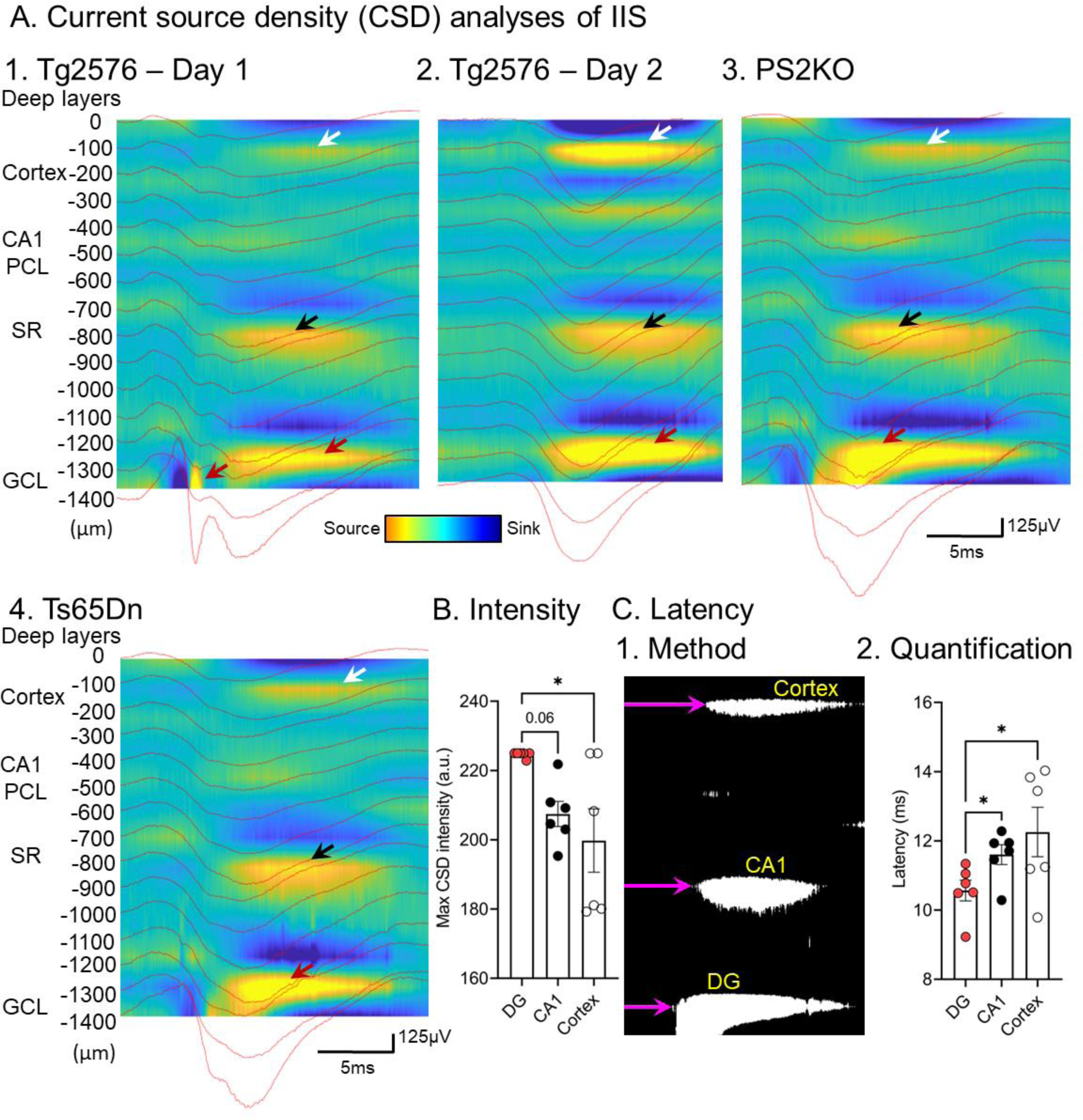
Current source density plots showing strong sources in the DG and additional sources in CA1 and cortex. (A) 1. A CSD plot is shown for an IIS recorded in a Tg2576 mouse. There were two current sources (yellow; red arrows) near the GCL and two additional sources were in area CA1 SR (black arrow) and cortex (white arrow). The sources in SR and cortex were relatively weak compared to GCL, in most cases. The depth is shown in 100μm increments corresponding to the distance between electrodes; the zero point corresponds to the first electrode in the deep cortical layers. Sources are denoted by warm colors (yellow/orange) and sinks by blue shades. 2. Same mouse as in 1 but the CSD is shown for an IIS recorded on a different day (Day 2). Note the robust GCL source (red arrow) is similar to Day 1 and therefore reproducible. The sources in SR (black arrow) and cortex (white arrow) were also reproducible. 3. Same as in 1-2 but the recordings were from a PS2KO mouse. Note a prominent source in the DG (red arrow) and additional sources in SR (black arrow) and cortex (white arrow). 4. Same as 3 but the recordings were from a Ts65Dn mouse. There were sources in the DG, SR and cortex as for other mouse lines. (B) Quantification of maximal CSD intensity of current sources in the DG, SR and cortex. The max CSD intensity was significantly different between the DG, CA1 SR and cortex (Friedman test (3)=8.09, p=0.009). *Post-hoc* comparisons showed a trend towards higher CSD intensity in the DG vs. CA1 (p=0.06) and a statistically significant difference between DG and cortex (p=0.01). Note that when IIS was used as the sample size unit, rather than mice, the DG was significantly greater than both CA1 and cortex (see text). (C) 1. Methods used to quantify latency to DG, CA1 SR and cortical source in the CSD. CSD plots were rendered black and white, and a threshold was determined based on the minimum CSD intensity quantified in B. The intensity was further reduced by 2 SEM. to adequately capture even the weakest current sources such as those in cortex (see Methods). In white are shown current sources that exceeded the threshold. Purple arrows show the latency to each source relative to the onset of each CSD. 2. We found a statistically significant difference in the latency to DG, CA1 SR and cortical current source (repeated measures ANOVA, F(1.17, 5.83)=6.9, p=0.03). *Post-hoc* comparisons showed that current sources in the DG were significantly earlier compared to CA1 SR (p=0.007) or cortex (p=0.001).

Notably, the majority of CSDs showed an early source in the DG (Fig 8A1-4) so we next analyzed the latency to each source. We used a threshold that allowed us to visualize even the weakest current sources (such as those in cortex; Fig 8C1) which was based on the lowest maximal CSD intensity (see Methods). We found a statistically significant difference in the latency to the onset of each current source (repeated measures ANOVA, F(1.17, 5.83)=6.9, p=0.03; Fig 8C2). *Post-hoc* comparisons showed that current sources in the DG were significantly earlier compared to CA1 SR (p=0.007; Fig 8C2) or cortex (p=0.001; Fig 8C2).

These data suggest that the DG could initiate the IIS and additional areas follow. A simple explanation would be that the DG triggers activity in CA1 by the trisynaptic pathway, and then cortical areas are activated after CA1 excitation of the entorhinal cortex. The synaptic delays would be milliseconds, consistent with the LFPs that have millisecond delays. However, when decompressed, the onsets of the LFPs are hard to define and it is not always clear the DG LFP begins first. These limitations are raised in the Discussion.

### VII. Selective chemogenetic inhibition of MS cholinergic neurons significantly reduced IIS

IIS occurred during REM sleep, a sleep period when MS cholinergic tone is markedly increased compared to SWS or wakefulness (Jasper and Tessier, 1971; Vazquez and Baghdoyan, 2001). Therefore we asked whether IIS rate would be affected if we silenced MS cholinergic neurons selectively. To test this hypothesis, we used Tg2576 transgenic mice because they show significantly more IIS compared to PS2KO or Ts65Dn transgenic mice (see Fig 2F, 3B3, 3C3).

Tg2576 transgenic mice were crossed to ChAT-Cre homozygous mice so that offspring that were positive for the Tg2576 transgene and positive for Cre (ChAT-Cre::Tg2576) would selectively express Cre recombinase in MS cholinergic neurons (see Methods). Next, ChAT-Cre::Tg2576 mice were injected with AAV5-hSyn-DIO-hM4D(Gi)-mCherry in MS and a depth electrode was implanted in the DG (Fig 9A). Next, we asked whether selective chemogenetic silencing of MS cholinergic neurons would be sufficient to reduce the number of IIS. Thus, we used a chemogenetic approach (Fig 9B) where ChAT-Cre::Tg2576 mice were recorded for a 2 hr-long baseline period and the number of IIS per hr was determined. Next, the mice were injected i.p. with a 3 mg/kg dose of CNO. Immediately after a 30 min waiting period for CNO to peak in the brain (Jendryka et al., 2019), recordings were made for the next 1-2 hr. The number of IIS per hr was determined for the first 1 hr to judge the effect of CNO, and the second 1 hr was used to determine if IIS rate returned to baseline. We also conducted 2 control experiments (Fig 9C-D). Thus, control experiments used either saline injection in ChAT-Cre::Tg2576 mice (Control #1; Fig 9C) or CNO in Tg2576 mice that were not injected with AAV (Control #2; Fig 9D).

**Figure 9:**
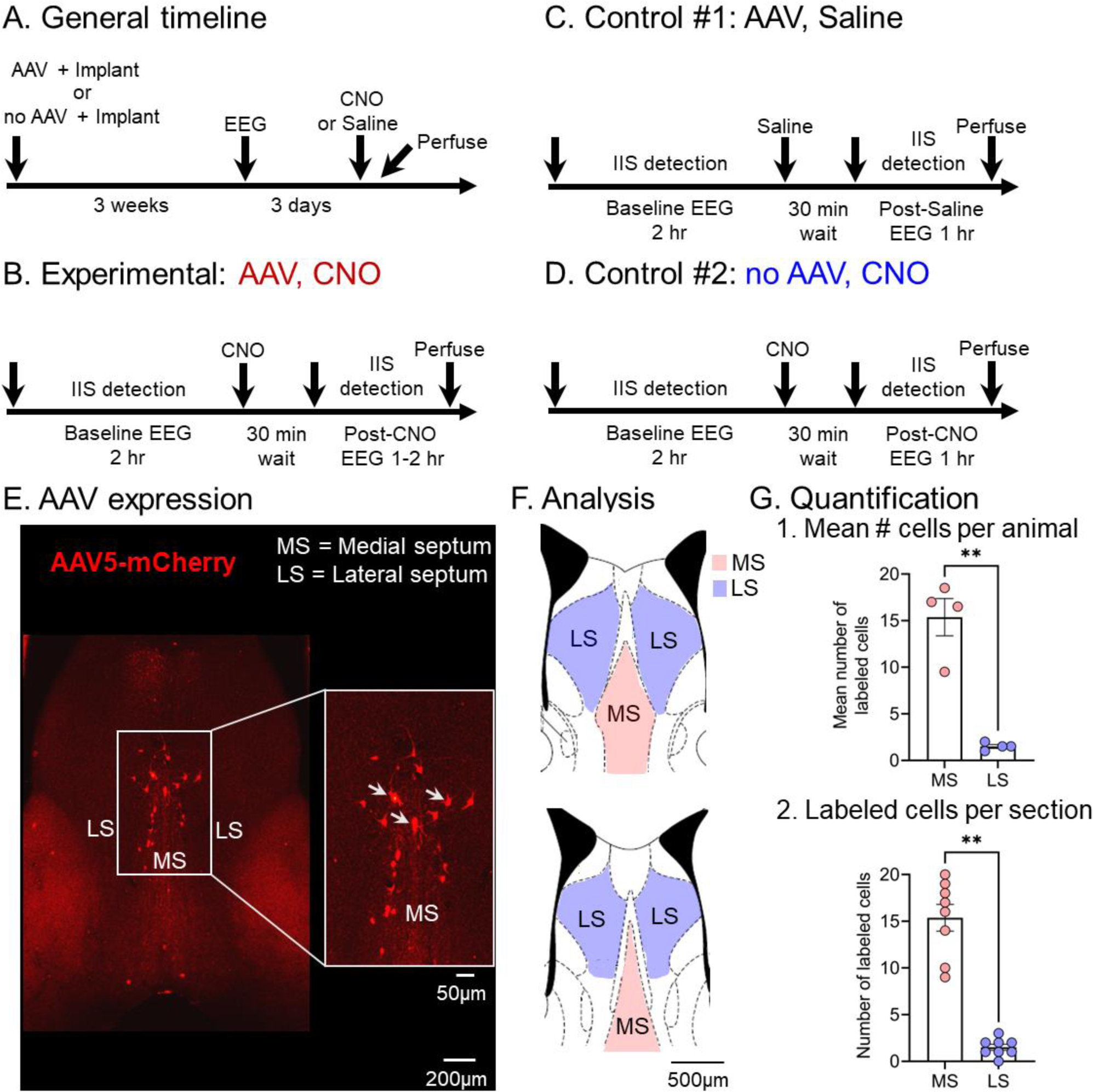
Experimental approach for chemogenetics and confirmation of viral expression in MS. (A) Experimental approach. ChAT-Cre::Tg2576 mice were injected with AAV5-hSyn-DIO-hM4D(Gi)-mCherry (AAV) in MS and in the same surgery mice were implanted with an electrode in the DG. Mice received CNO or saline. An additional cohort of Tg2576 mice were implanted and received CNO but no AAV was injected to serve as additional controls (no AAV; see below). After 3 weeks to allow for expression of the virus and recovery from surgery, mice were video-EEG recorded continuously for 3 consecutive days and then CNO or saline was injected. Afterwards, mice were euthanized to confirm viral expression (see Methods). (B) Timeline of the CNO experiments using ChAT-Cre::Tg2576 mice injected with AAV5-hSyn-DIO-hM4D(Gi)-mCherry (AAV, CNO). A 2 hr-long baseline EEG was recorded and then 3 mg/kg CNO was injected i.p. After waiting for 30 min, we counted IIS for the next 2 hr and afterwards the mice were euthanized. (C) Same as in B but for a different cohort of ChAT-Cre::Tg2576 mice injected with AAV5-hSyn-DIO-hM4D(Gi)-mCherry where saline was injected instead of CNO (AAV, Saline). IIS rate was quantified for 1 hr following the 30 min waiting period. (D) Same as in C but Tg2576 mice were used. These mice were not injected with AAV but they were injected with CNO (no AAV, CNO). (E) Representative viral expression of AAV5-hSyn-DIO-hM4D(Gi)-mCherry in MS of a ChAT-Cre::Tg2576 mouse. Inset shows MS at higher magnification and white arrows point to individual cells expressing AAV5-hSyn-DIO-hM4D(Gi)-mCherry. MS = Medial septum, LS = Lateral septum. (F) Schematics of the brain sections used for analysis are shown. Top, a section corresponding to approximately +0.6 mm A-P from Bregma. Bottom, a section corresponding to approximately + 0.8 mm A-P (adapted from Franklin and Paxinos, 1997). (G) 1. The mean number of mCherry-labeled cells per animal was significantly greater in MS vs. LS (paired t-test, t*_crit_*=7.44, p=0.005; n=4 mice). 2. There were significantly more mCherry-labeled cells in MS vs. LS when comparisons were made using the number of cells per section instead of the mean per animal (Wilcoxon signed rank test, p=0.008; n=8 sections from 4 mice).

We assessed AAV5-hSyn-DIO-hM4D(Gi)-mCherry expression in MS using the fluorescent tag mCherry (see Methods) and a representative example is shown in Fig 9E. To address the extent of mCherry expression and how restricted it was to the MS we quantified mCherry-labeled cells in the MS and lateral septum (LS) using 2 sections per mouse (Fig 9F). The data showed that mCherry expression was largely restricted to MS and relatively rare in LS. When the mean of the 2 sections was used for comparison of mice, the differences were significant (paired t-test, t*_crit_*=7.44, p=0.005; n=4 mice; Fig 9G1). When sections were compared, differences were also significant (Wilcoxon signed rank test, p=0.008; n=8 sections; Fig 9G2).

Fig 10A shows a representative baseline recording. Note that IIS occurred when theta/delta ratio was above the threshold to define REM sleep (dotted line in Fig 10A-B). Within the 1 hr after CNO injection, the number of IIS occurring during REM sleep was markedly reduced (Fig 10A vs. 10B) compared to baseline (paired t-test, t*_crit_*=4.59, p=0.01, n=4 mice; Fig 10C). This cohort of mice formed the “CNO group.”

**Figure 10:**
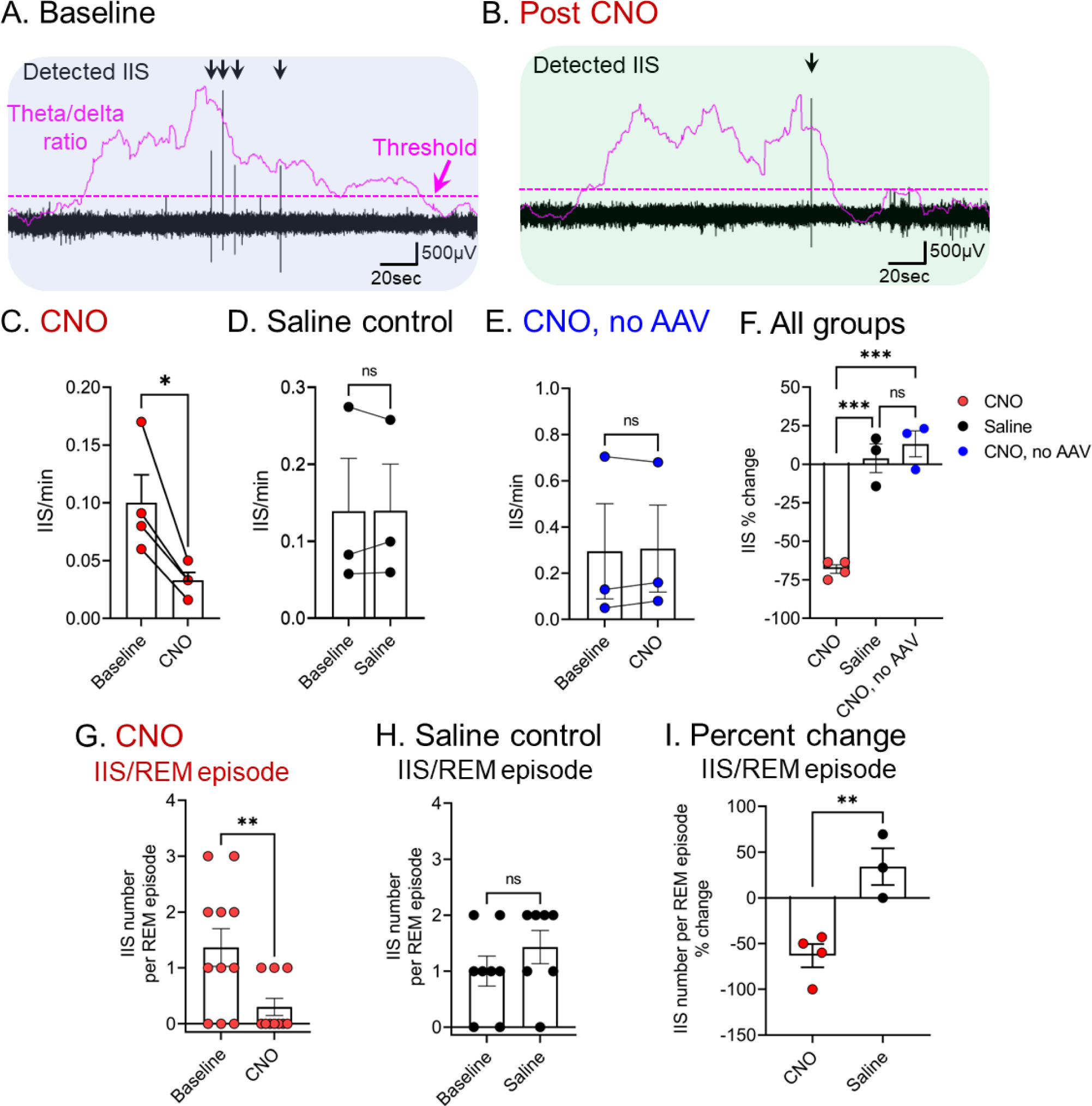
IIS rate is significantly reduced by selective chemogenetic inhibition of MS cholinergic neurons. (A) Representative recordings during a baseline. Theta/delta ratio is shown in purple shading and the threshold is marked by a purple dotted line; detected IIS are noted with a black arrow. Note that IIS occurred during a period of increased theta/delta ratio. (B) In the same mouse as A, CNO was injected, and the recording is shown after waiting 30 min for CNO to peak in the brain. Note the reduced number of IIS. (C) IIS rate was reduced 1 hr vs. baseline (paired t-test, t*_crit_*=4.59, p=0.01). (D) Same as in C but saline was injected instead of CNO. IIS rate did not significantly change between baseline and after saline injection (paired t-test, *t*_crit_=0.007, p=0.95, n=3 mice) (E) IIS rate in Tg2576 mice without AAV was not significantly affected by CNO injection (paired t-test, *t*_crit_=0.31, p=0.78, n=3 mice). (F) Percent change in IIS rate between groups (CNO; Saline control; CNO no AAV). One-way ANOVA showed that the percent change in IIS rate was significantly different (F(2, 7)= 47.95, p<0.0001) between the 3 groups. *Post-hoc* comparisons revealed a significantly greater reduction in IIS rate in the CNO group (68.1±2.8%, n=4 mice) compared to controls (saline injection: p=0.0003, n=3 mice; CNO no AAV; p=0.0002, n=3 mice). No statistically significant differences were found between control groups (p=0.76). (G) IIS per REM sleep episode in the CNO group. This group was Chat-Cre::Tg2576 and received AAV and CNO. The number of IIS per REM sleep episode was significantly reduced between baseline and after CNO injection. Comparisons were based on all REM episodes (Wilcoxon signed rank test, p=0.02, n=4 mice). (H) IIS per REM sleep episode in the Saline control group. This group was Chat-Cre::Tg2576 and received AAV and saline instead of CNO. The number of IIS per REM episode did not significantly change (Wilcoxon signed rank test, p=0.31, n=3 mice) between baseline and after saline injection. As in G, comparisons were based on all REM episodes. Thus, Saline controls lacked effects observed in the CNO group. (I) Percent change in the mean number of IIS/REM episode per animal showed a statistically significant difference between the CNO and Saline group (unpaired t-test, *t*_crit_=4.31, p=0.007, CNO, n=4 mice; Saline, n=3 mice).

We also used 2 control groups. The first control experiment used saline injection in ChAT-Cre::Tg2576 mice injected with AAV5-hSyn-DIO-Hm4D(Gi)-mCherry (Saline control). There were no statistically significant differences in IIS rate between baseline and after saline injection (paired t-test, t*_crit_*=0.007, p=0.95, n=3 mice; Fig 10D). The second control experiment used CNO injection in Tg2576 transgenic mice without AAV5-hSyn-DIO-hM4D(Gi)-mCherry (“CNO, no AAV group”). There were no differences in IIS rate between baseline and after CNO injection (paired t-test, t*_crit_*=0.31, p=0.78, n=3 mice; Fig 10E).

The percent changes in IIS rate were significantly different between the 3 groups (CNO; Saline control; CNO no AAV; one-way ANOVA, F(2, 7)= 47.95, p<0.0001; Fig 10F). *Post-hoc* comparisons revealed a significantly greater reduction in IIS rate in the CNO group (68.1±2.8%, n=4 mice) compared to controls (Saline control: p=0.0003, n=3 mice; CNO no AAV; p=0.0002, n=3 mice), but not between the 2 control groups (p=0.76; Fig 10F).

Additional analyses examined IIS per REM sleep episode. The number of IIS per REM sleep episode was lower after CNO injection compared to baseline (Wilcoxon signed rank test, p=0.02; Fig 10G). In contrast, the number of IIS per REM episode did not significantly change after saline injection (Wilcoxon signed rank test, p=0.31; Fig 10H). We next compared percent change in the mean number of IIS/REM episode per animal and found a statistically significant difference between the CNO and Saline group (unpaired t-test, t*_crit_*=4.31, p=0.007; Fig 10I) similar to the effects shown for IIS rate in Fig 10F.

### VIII. MS cholinergic silencing did not significantly change REM sleep duration, number of REM bouts, theta power during REM or latency to REM

We next asked whether the reduction in IIS rate could be explained by any effects on REM sleep characteristics such as duration, number of bouts, or theta power during REM. We found no statistically significant differences in total duration of REM sleep between baseline and 1 hr post-injection either in mice injected with CNO (Wilcoxon signed rank test, p=0.87; Fig 11A1) or saline (paired t-test, t*_crit_*=0.13, p=0.90; Fig 11A2). Similarly, we did not find any statistically significant differences in the number of REM bouts in the CNO (paired t-test, t*_crit_*=0.52, p=0.63; Fig 11B1) and the Saline-injected controls (Wilcoxon signed rank test, p>0.99; Fig 11B2). The duration of REM bouts did not significantly change either in the CNO-injected mice (Wilcoxon signed rank test, p=0.91; Fig 11C1) or Saline-injected controls (paired t-test, t*_crit_*=0.28, p=0.78; Fig 11C2). We also quantified theta power during REM. Notably, theta power during REM did not significantly change between baseline and after injection either in mice injected with CNO (paired t-test, t*_crit_*=0.62, p=0.57; Fig 11D1) or saline (paired t-test, t*_crit_*=2.06, p=0.17; Fig 11D2). Indeed, the percent change in theta power between baseline and after injection was not significantly different between CNO and Saline-injected mice (paired t-test, t*_crit_*=2.64, p=0.12; Fig 11E). These data suggest that inhibition of IIS by MS silencing was not due to a reduction in REM or theta.

**Figure 11:**
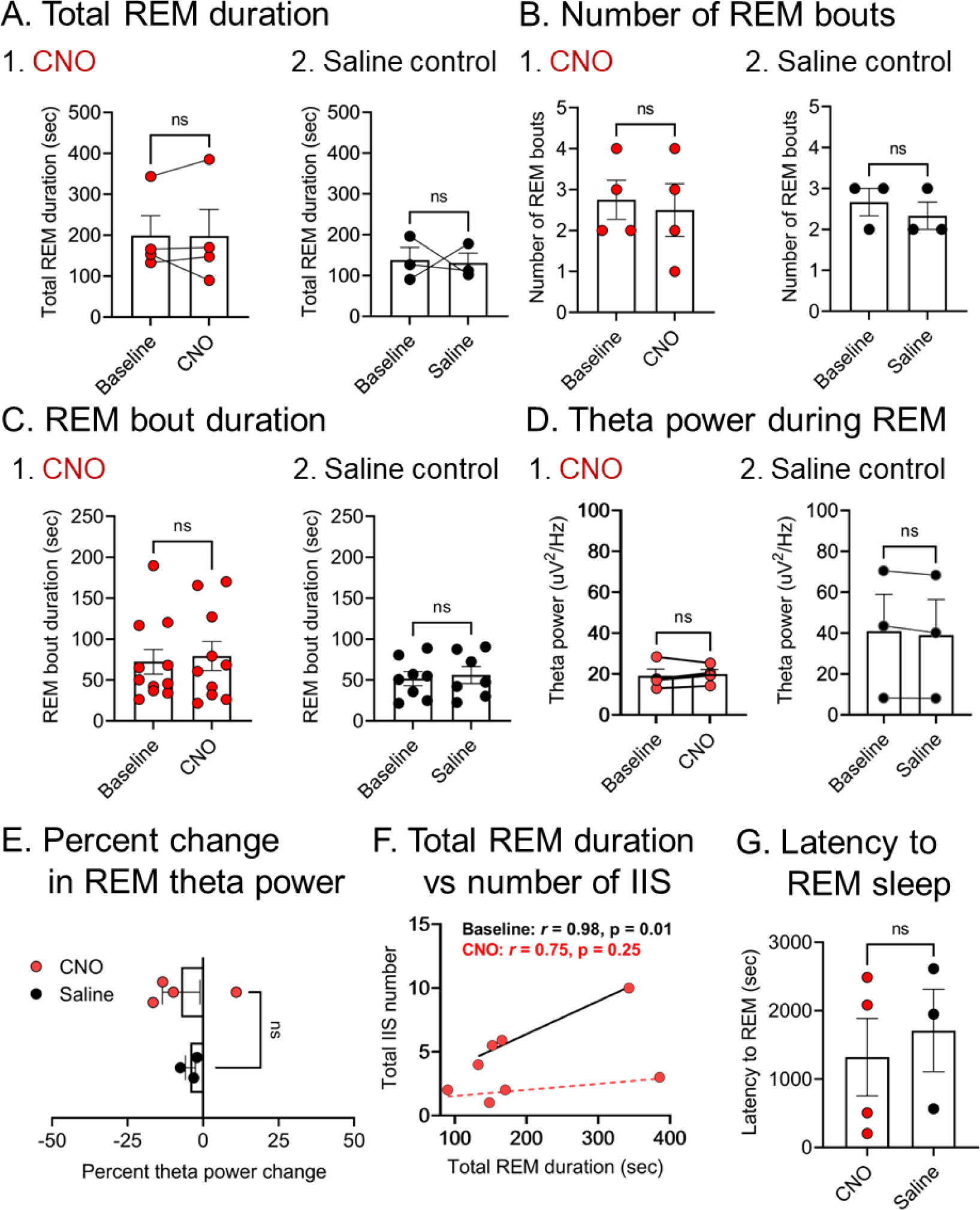
REM sleep characteristics are not compromised by MS cholinergic silencing. (A) Total REM duration. 1. The total duration of REM sleep during the 60 min (1 hr) of the baseline and 30-90 min (1 hr) after CNO injection is shown. We found no statistically significant differences between baseline and post-CNO (Wilcoxon signed rank test, p=0.87). 2. Same as in 1 but for Saline controls. There were no significant differences (paired t-test, *t*_crit_=0.13, p=0.90). (B) Number of REM bouts. 1. The number of REM sleep bouts is shown for mice injected with CNO. We found no differences between baseline and after CNO injection (paired t-test, *t*_crit_=0.52, p=0.63). 2. Same as in 1 for Saline-injected controls. We found no statistically significant differences (Wilcoxon signed rank test, p>0.99). (C) REM bout duration. 1. We found no statistically significant differences for REM bout duration between baseline and 1 hr post-CNO injection (Wilcoxon signed rank test, p=0.91). 2. Same as in 1 but for Saline-injected mice. No statistically significant differences were found (paired t-test, *t*_crit_=0.28, p=0.78). (D) Theta power during REM sleep. 1. We found no statistically significant differences in theta power between baseline and 1 hr post-CNO injection (paired t-test, *t*_crit_=0.62, p=0.57). 2. Same as in 1 but for Saline-injected controls. We found no differences (paired t-test, *t*_crit_=2.06, p=0.17). (E) Percent change in theta power. The percent change in theta power between baseline and after injection was not significantly different between CNO and Saline-injected mice (paired t-test, *t*_crit_=2.64, p=0.12). (F) Correlation analyses between total REM duration and the total number of IIS in REM. We found a statistically significant positive correlation between the total duration of REM sleep and the total number of IIS during baseline (Pearson *r* = 0.98, p=0.01) but not during the 1 hr after CNO (Pearson *r* = 0.75, p=0.25), n=4 mice. (G) Latency to REM sleep. We found that the latency to REM sleep was not significantly different between CNO and Saline-injected controls (unpaired t-test, t*_crit_*=0.46, p=0.66).

In an additional analysis, we compared whether the total duration of REM sleep would correlate with the total number of IIS we recorded. Indeed, we found a statistically significant positive correlation between the total duration of REM and number of IIS during baseline (Pearson *r* = 0.98, p=0.01; Fig 11F1) but not after CNO (Pearson *r* = 0.75, p=0.25; Fig 11F1), presumably due to the marked reduction in IIS. Last, we quantified the latency to first REM sleep episode and asked whether it was affected by CNO injection. Notably, CNO injection did not change the latency to the first REM sleep episode compared to Saline-injected controls (unpaired t-test, t*_crit_*=0.46, p=0.66). This is important because a longer delay to REM sleep may contribute to sleep disturbances and that has been proposed as a diagnostic biomarker for AD (Bliwise et al., 1989; D’Rozario et al., 2020; Zhang et al., 2022).

### IX. Lack of rebound in IIS rate and no “ectopic” occurrence of IIS following MS cholinergic silencing

We next asked whether chemogenetic silencing of MS led to any side effects. We hypothesized that potential side-effects could be a rapid increase in IIS rate after MS silencing and/or “ectopic” occurrence of IIS in behavioral states besides REM sleep.

We found that IIS rate was significantly different between baseline, 1 hr post-CNO and 2 hr post-CNO (Mixed-effects model, F(2, 5)= 8.73, p=0.02; Fig 12A-B1). *Post-hoc* comparisons showed that IIS rate was significantly reduced during the 30-90 min (first hr) after CNO injection (p=0.03) compared to baseline and returned to baseline levels during the 90-150 min (second hr) post-CNO (baseline vs. 2 hr; p=0.41). The direction of the IIS rate during the first hr was a reduction and it switched to an increase during the second hr; quantified as the percent change in IIS rate, the differences were statistically significant (paired t-test, *t*_crit_=8.01, p=0.01; Fig 12B2). Thus, IIS rates decreased within 1 hr while they increased within 2 hr post-CNO, returning to a rate that was not significantly different from baseline. In summary, there was no evidence of rebound.

**Figure 12:**
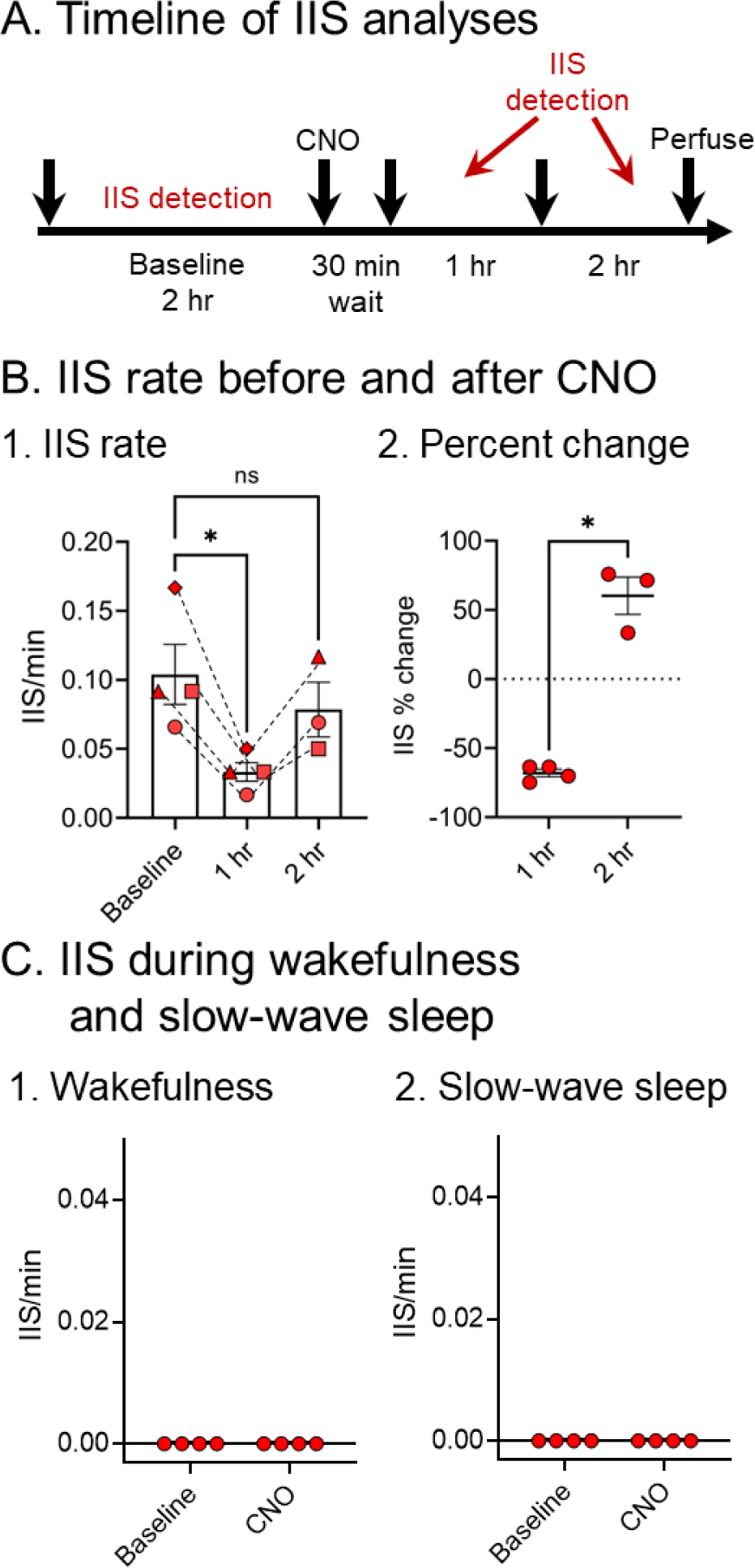
Chemogenetic silencing of MS did not induce a rebound increase in IIS rate or ectopic occurrence of IIS. (A) Timeline of IIS analyses. IIS rate was quantified for 1 hr during the baseline, 30-90 min (first hr) after CNO injection and 90-150 min (second hr) post-CNO. (B) Changes in IIS rate following CNO injection in the CNO group. These animals were ChaT-Cre::Tg2576 mice injected with AAV. 1. IIS rate was significantly different between the baseline, the first hr post-CNO and the second hr post-CNO (Mixed-effects model, F(2, 5)= 8.73, p=0.02; Fig 7A1). *Post-hoc* comparisons showed a statistically significant difference between baseline and the first hr post-CNO (p=0.03). There was no difference in IIS rate between baseline and the second hr post-CNO (p=0.41), suggesting a recovery of IIS rate. 2. The percent change in IIS rate between the first and the second hr post-CNO was significantly different (paired t-test, *t*_crit_=8.01, p=0.01). The direction switched from a decrease during the first hr post-CNO to an increase during the second hr post-CNO (relative to baseline). (C) Occurrence of IIS during wakefulness and SWS during the first hr after CNO injection. 1. IIS were not detected during wakefulness. 2. Similar to wakefulness, IIS were not detected during SWS.

We also checked whether IIS during the first 1 hr occurred during behavioral states besides REM because these data would suggest that there was “ectopic” occurrence of IIS. We did not detect any IIS either during wakefulness (Fig 12C1) or SWS (Fig 12C2). Therefore, MS silencing did not appear to cause “ectopic” IIS.

## DISCUSSION

### Summary of main findings

The goal of this study was to understand sites in the brain where IIS are robust and the role of MS cholinergic neurons in controlling IIS. We found that IIS is a robust abnormality in 3 mouse lines that simulate features of AD. We also found that IIS occurred primarily in REM sleep in all 3 mouse lines. Using chemogenetics, we confirmed that selective silencing of MS cholinergic neurons reduced IIS rate. In addition, multi-channel LFP recordings revealed that IIS amplitude was maximal in the GCL relative to area CA1 and overlying cortex. Our findings are surprising in light of the studies showing that most granule cells (GCs) are relatively quiet in normal mice and rats (Jung and McNaughton, 1993; Diamantaki et al., 2016; GoodSmith et al., 2017; Senzai and Buzsáki, 2017; Pofahl et al., 2021). We also provide novel evidence for a contributing role of MS cholinergic neurons in IIS generation which may serve as a new strategy to abate IIS. This idea is also surprising given the widespread notion that the cholinergic system in AD should always be enhanced, not depressed (Lahiri and Farlow, 2003; Trinh et al., 2003). Thus, our data suggest an unexpected way to counteract IIS early in the disease.

### IIS represent a common EEG disturbance in different mouse lines

We found that IIS occurred in all transgenic mice but not in the controls. Although IIS are known to occur in Tg2576 transgenic mice (Kam et al., 2016; Lisgaras and Scharfman, 2022a), comparative recordings of different brain areas have not been made. Moreover, IIS in PS2KO and Ts65Dn transgenic mice have not been studied. Therefore, our findings are novel. Although novel, the results are consistent with past studies. For example, PS2KO mice have increased excitability, manifested by a lower seizure threshold when seizures are triggered using the kindling paradigm (Beckman et al., 2020). However, spontaneous epileptiform activity has not been reported to date. It is possible that IIS contribute to higher seizure susceptibility of these mice. However, IIS in epilepsy are not always considered to promote seizures (Avoli et al., 2006; Staley and Dudek, 2006; Karoly et al., 2016). One additional indication of hyperexcitability in PS2KO mice is HFOs, which we reported recently (Lisgaras and Scharfman, 2022a). Thus, PS2KO mice show two types of spontaneous abnormalities in the EEG, IIS and HFOs, supporting the view that the mice have hyperexcitability. That is interesting in light of the clinical findings of *PS2* mutations in AD, where hyperexcitability has been reported (Jayadev et al., 2010c; Zarea et al., 2016; Lehmann et al., 2021).

The Ts65Dn transgenic mice we recorded also showed robust IIS which has not been reported before. Taken together, the presence of robust IIS in 3 mouse lines with different causal mechanisms suggests that IIS can arise for multiple reasons, consistent with epilepsy where IIS occur in diverse types of epilepsy. It remains to be determined whether IIS in epilepsy and AD share similar characteristics and whether they occur in similar brain areas.

Our results also suggest that IIS could be a promising EEG signature to consider as an outcome measure in AD. Notably, the occurrence of IIS at a range of different ages argues against the possibility that IIS is a temporary EEG manifestation. Thus, IIS could be considered as a biomarker in AD, as noted elsewhere (Kam et al., 2016; Lehmann et al., 2021).

### IIS were most frequent in Tg2576 transgenic mice

Among all 3 mouse lines we examined, we found that IIS were most frequent in Tg2576 transgenic mice (vs. PS2KO or Ts65Dn mice). HFOs were also more frequent in Tg2576 mice vs. PS2KO or Ts65Dn mice (Lisgaras and Scharfman, 2022a). Higher rates of IIS and HFOs in Tg2576 mice could be due to the elevated levels of Aβ in Tg2576 mice (Hsiao et al., 1996) which do not occur in PS2KO (Herreman et al., 1999) and Ts65Dn mice (Salehi et al., 2006). This idea is consistent with a study showing increased calcium transients in neurons around plaques, suggesting that there is hyperexcitability around plaques (Busche et al., 2008) However, IIS and HFOs can occur before plaques are detected (Kam et al., 2016; Lisgaras and Scharfman, 2022a) so elevated levels of oligomeric Aβ might play a role in hyperexcitability. Additional mechanisms may also be involved.

Tg2576 transgenic mice are also known to experience seizures (Bezzina et al., 2015; Kam et al., 2016; Lisgaras and Scharfman, 2022a). Thus, frequent IIS in Tg2576 transgenic mice may be caused or contribute to seizures in this mouse line. That idea is consistent with a previous study using APP/PS1 transgenic mice which showed that IIS rate correlated with a higher seizure burden (Gureviciene et al., 2019).

### IIS during REM sleep and cholinergic control by medial septum

We found that IIS in the 3 mouse lines we studied were robust in REM sleep. Acetylcholine (ACh) levels during REM sleep are increased to a greater extent compared to other behavioral states such as wakefulness or SWS (Jasper and Tessier, 1971; Vazquez and Baghdoyan, 2001). Thus, it is possible that increased ACh levels during REM sleep may facilitate the occurrence of IIS. Indeed, cholinergic facilitation of IIS and seizures is known to occur in epilepsy (Friedman et al., 2007; Maslarova et al., 2013; Mikroulis et al., 2018). Consistent with these data, our previous study (Kam et al., 2016) found that i.p. injection of the cholinergic antagonist atropine reduced IIS on REM as well as NREM (which primarily includes SWS). However, the reduction of IIS in REM by atropine was confounded by a shortening of REM sleep duration after atropine administration. Also, the study (Kam et al., 2016) did not provide specificity for the involvement of MS cholinergic neurons as the current study.

The prior study using atropine prompted us to implement an approach that would minimize effects on REM such as those related to reduced REM sleep duration. Thus, we chose a chemogenetic approach to interfere with cholinergic neurotransmission selectively by targeting MS cholinergic neurons. Using this selective approach, we found that the number of IIS was significantly reduced without any significant effects in REM sleep duration or the number and duration of REM sleep bouts. These results support a role of MS cholinergic neurons in IIS generation.

Our data contrast with the widespread notion that cholinergic function should always be increased in AD (Hampel et al., 2018). This dominant view comes from observations of robust basal forebrain neurodegeneration which typically occurs late in AD progression (Bartus et al., 1982; Hampel et al., 2018; Kim et al., 2019; Falangola et al., 2021). Thus, the late degeneration of MS cholinergic neurons makes it appear that ACh levels are always too low in AD, and need to be increased. However, it has been shown that cholinergic markers like ChAT are high in early AD stages like MCI (Hyman et al., 1987; DeKosky et al., 2002; Mufson et al., 2008). In addition, increasing ACh in late stages of AD using cholinesterase inhibitors like donezepil has not been highly effective (Lahiri and Farlow, 2003; Trinh et al., 2003). In our view, cholinergic activity may be elevated early and decay late in AD. Thus, silencing hyperactive MS cholinergic neurons in early AD may be beneficial. Silencing MS cholinergic neurons may also protect them from excessive activation and consequent deterioration.

It is also important to consider that other cholinergic nuclei may also control IIS. For instance, cholinergic nuclei located in brainstem may be important as they appear to contribute to REM sleep (Van Dort et al., 2015). In addition, other systems as well as neuropeptides that participate in sleep-wake transitions should also be considered (Saper et al., 2001; Taheri et al., 2002).

### Brain state dependence of IIS in AD: humans vs. mice

Our data suggesting that IIS in 3 mouse lines predominantly occur in REM sleep, may appear at odds with human AD/MCI literature suggesting that epileptiform activity only is prevalent in NREM sleep (Vossel et al., 2016; Lam et al., 2017). However, review of the data in the literature suggests that it may not always be the case that IIS in human AD/MCI hardly ever occur during REM sleep.

One of the reasons to be cautious is that many studies either did not analyze REM (at least initially) (Lam et al., 2017) or did not distinguish between REM and NREM sleep (Vossel et al., 2016; Lam et al., 2017). Indeed, in a re-analysis (Brown et al., 2018) of the landmark foramen ovale study (Lam et al., 2017) where IIS were recorded with an electrode penetrating the foramen ovale, NREM sleep was separated from REM and IIS were detected in REM, but at a lower rate than IIS in NREM. A prior scalp EEG study (Vossel et al., 2016) in humans with AD studied epileptiform activity in sleep stages ≥2. Sleep stages ≥2 include exclusively NREM sleep and exclude REM sleep. Consequently, a lack of distinction between different sleep stages may have given the impression that IIS in humans with AD/MCI do not occur in REM. Notably, in Tg2576 mice there is a predominance of IIS in REM as the present study shows, but one still can record many IIS in NREM, especially at ages over 7 months (Kam et al., 2016). Nevertheless, more studies are needed to conclusively address brain state dependence of IIS in humans and animals.

Another important consideration is that humans especially at an advanced AD stage may show reduced REM duration (Vitiello et al., 1984). This makes it harder to show that IIS occur in REM. Indeed, one individual that was recorded with foramen ovale electrodes (Lam et al., 2017) at an advanced AD stage recorded no REM sleep during the time interval of the recording (Brown et al., 2018). Notably, a similar recording in one MCI patient that captured REM sleep did indeed record IIS during REM (Brown et al., 2018). Thus, the duration of EEG monitoring is a critical factor in reliably assessing whether epileptiform activity occurred in different behavioral states. Indeed, serial EEGs (≥2) or long-term video EEG monitoring (≥24 hr) are more effective at detecting epileptiform activity compared to routine EEGs (Vossel et al., 2013). Also, using receiver operating characteristics analyses, the sensitivity of detecting epileptiform activity in AD is significantly increased (Horváth et al., 2017) similar to epilepsy (Doppelbauer et al., 1993; Faulkner et al., 2012). Thus, routine EEGs might have generally underestimated IIS and specifically IIS in REM because REM is typically shorter compared to NREM (Liedorp et al., 2010).

Another important insight comes from a study looking at IIS in AD patients with and without epilepsy as well as healthy controls (Lam et al., 2020). The study showed that IIS were more frequent during REM and wakefulness in those AD patients that have diagnosed epilepsy (vs. those that were not diagnosed with epilepsy). In addition, IIS during REM and wakefulness showed both good specificity (85.7%) and sensitivity (85.7%) in predicting clinical seizures. Interestingly these results suggesting a close relationship between IIS and seizures are consistent with the use of IIS during REM to localize epileptogenic foci (Sammaritano et al., 1991).

To conclude, we propose a re-examination of the idea that IIS only occur in NREM by implementing longer or repeated EEG recording and sleep staging that differentiates between NREM and REM.

### The DG as a possible source of IIS

Excitability in the DG is known to change early during AD progression. For example, in amnestic MCI, fMRI showed high activity in the DG (Bakker et al., 2012). In mice, markers of neuronal activity such as c-Fos and ΔFosB were elevated in the J20 mouse line in early life (Corbett et al., 2017). Reduced calbindin-D28K (Palop et al., 2003) and increased neuropeptide Y (Palop et al., 2007) in GCs of J20 mice are also consistent with hyperactive GCs. In vitro, slices of Tg2576 mice at 2-3 months of age showed increased excitability of GCs (Alcantara-Gonzalez et al., 2021). In APP/PS1 mice, GCs were depolarized relative to controls (Jiang et al., 2021). In vivo, we showed HFOs in the DG at just 1 month of age (Lisgaras and Scharfman, 2022a).

Our study is the first to show that the DG dominates the activity that occurs during IIS, at least along the cortical-CA1-DG axis. We showed that in all 3 mouse lines we tested IIS appeared to be maximal in amplitude in the GCL and accompanied by strong current sources. A larger IIS amplitude would suggest that a greater number of neurons fire to produce an IIS compared to an IIS that is characterized by a smaller amplitude. Indeed, multiunit activity during IIS in epilepsy is found in brain areas where IIS are likely to be generated (Ulbert et al., 2004; Guth et al., 2021). Overall, these results suggest that the DG may contribute to a larger extent to IIS compared to either area CA1 or overlying cortex where IIS appear to occur with a significantly lower amplitude. Notably, we also detected strong current sources in the DG. The current sources in the DG preceded those in CA1 and cortex suggesting that DG currents are an early contributor to IIS.

### Relationship of IIS to dentate spikes

In normal rodents, a large spike occurs spontaneously in the DG EEG, called a dentate spike (Bragin et al., 1995; Buzsáki et al., 2003; Headley et al., 2017; Dvorak et al., 2021). It is interesting to consider the possible relationship of IIS to dentate spikes in light of the large IIS in the DG in the three mouse lines we studied. On the one hand, IIS are distinct because they are recorded throughout many brain regions, but the dentate spike is more localized (Bragin et al., 1995; Buzsáki et al., 2003; Headley et al., 2017; Dvorak et al., 2021). Moreover, previous studies suggest the entorhinal cortex drives dentate spikes (Bragin et al., 1995; Dvorak et al., 2021) whereas our CSDs showed that a prominent DG source of the IIS was in the inner molecular layer as well as additional sources in CA1 and cortex. Nevertheless, it is possible that dentate spikes occur early in life and then progressively recruit other brain regions and thus begin to have triggers other than the entorhinal cortex as pathology in the mice develop.

### Limitations and future directions

The data showing that MS silencing reduces IIS suggest that the MS could be targeted to alleviate from IIS-related memory impairments. Although cognitive tests were not studied here, they could be in the future. Notably, other investigators have reported that rats with IIS have impaired memory retrieval (Kleen et al., 2010). Also, other investigators have reduced hyperexcitability (although not necessarily IIS) with the anticonvulsant levetiracetam and found that cognition improved in those individuals with amnestic mild cognitive impairment (Bakker et al., 2012). In this context, a recent study found that the effect of levetiracetam was robust in those individuals with AD that showed epileptiform activity in the EEG (Vossel et al., 2021). It will also be valuable to determine if MS silencing could alleviate amyloid burden or reduce phosphorylated tau.

Other limitations include determining the exact location of electrodes. Therefore, we cannot discount that small differences in electrode location could have influenced the results. We also recognize some cohorts of mice were small. Therefore, we made conclusions based on what findings were consistent across all mice.

Also, in this study, we compared sites along the cortical-CA1-DG axis. However, we recognize that other areas of the brain may also show large IIS. It is also important to consider that IIS in the different mouse lines may not reflect the same electrophysiological phenomenon. Indeed, in epileptic animals, IIS may be due to different mechanisms, such as glutamatergic EPSPs underlying a paroxysmal depolarization shift (Matsumoto and Marsan, 1964; Ayala et al., 1973; Schwartzkroin and Prince, 1977; Johnston and Brown, 1981), or GABAergic mechanisms leading to synchronization of principal cells (Kokaia et al., 1994; Avoli et al., 1996; Muldoon et al., 2015). Future studies looking at mechanisms should also consider the effect of the genetic background on the results and note that IIS manifestation (such as IIS morphology) may vary with mouse line and age of the animal.

## CONCLUSIONS

Showing that the DG is a “hotspot” where IIS are consistently maximal and accompanied by early current sources is important as it suggests the DG as an important node in the pathophysiology (Ohm, 2007). Demonstration in 3 different mouse lines all with distinct characteristics of AD makes the findings stronger and likely to be important also. The idea that the DG has an important role in IIS is consistent with many observations of altered excitability in the DG in AD (Bakker et al., 2012) or mouse lines simulating features of AD (Palop et al., 2003; Palop et al., 2007; Corbett et al., 2017; Alcantara-Gonzalez et al., 2021). Indeed, this view is also strengthened by our previous study showing that the DG is an early contributor to increased hippocampal excitability as shown by the occurrence of DG HFOs *in vivo* (Lisgaras and Scharfman, 2022a). In addition, our findings pointing to the MS as a contributor to IIS activity is topical considering the urgent need for a better understanding of the role of subcortical structures in AD pathophysiology (Ehrenberg et al.). Future work targeting the DG as well as the MS is warranted as it may provide new avenues for therapeutic intervention in individuals at risk for developing AD.

## ACKNOWLEDGEMENTS

We would like to thank Dr. Ralph Nixon for providing the PS2KO mice and Dr. Stephen Ginsberg for providing the Ts65Dn mice. This project was supported by NIH grants R01 AG-055328 and R01 NS-106983, and the New York State Office of Mental Health.

## CONFLICTS OF INTEREST/ETHICAL PUBLICATION STATEMENT

None of the authors has any conflict of interest to disclose. We confirm that we have read the Journal’s position on issues involved in ethical publication and affirm that this report is consistent with those guidelines.

## DATA AVAILABILITY STATEMENT

Data will be available from the corresponding author upon reasonable request.

## Abbreviations

AD: Alzheimer’s disease
DG: Dentate gyrus
GCL: Granule cell layer
HFOs: High frequency oscillations
IIS: Interictal spike
MS: Medial septum
NREM: Non-rapid eye movement sleep
PCL: Pyramidal cell layer of CA1
PS2KO: Presenilin 2 knockout
REM: Rapid eye movement sleep
SR: Stratum radiatum of CA1
SWS: Slow-wave sleep

## REFERENCES

1. Alcantara-Gonzalez, D., Chartampila, E., Criscuolo, C., Scharfman, H. E., 2021. Early changes in synaptic and intrinsic properties of dentate gyrus granule cells in a mouse model of Alzheimer’s disease neuropathology and atypical effects of the cholinergic antagonist atropine. Neurobiol Dis. 152, 105274.

2. Apelt, J., Kumar, A., Schliebs, R., 2002. Impairment of cholinergic neurotransmission in adult and aged transgenic Tg2576 mouse brain expressing the swedish mutation of human β-amyloid precursor protein. Brain Res. 953, 17–30.

3. Avoli, M., Barbarosie, M., Lücke, A., Nagao, T., Lopantsev, V., Köhling, R., 1996. Synchronous GABA-mediated potentials and epileptiform discharges in the rat limbic system *in vitro*. J Neurosci. 16, 3912–3924.

4. Avoli, M., Biagini, G., de Curtis, M., 2006. Do interictal spikes sustain seizures and epileptogenesis? Epilepsy Curr. 6, 203–207.

5. Ayala, G. F., Dichter, M., Gumnit, R. J., Matsumoto, H., Spencer, W. A., 1973. Genesis of epileptic interictal spikes. New knowledge of cortical feedback systems suggests a neurophysiological explanation of brief paroxysms. Brain Res. 52, 1–17.

6. Bakker, A., Krauss, G. L., Albert, M. S., Speck, C. L., Jones, L. R., Stark, C. E., Yassa, M. A., Bassett, S. S., Shelton, A. L., Gallagher, M., 2012. Reduction of hippocampal hyperactivity improves cognition in amnestic mild cognitive impairment. Neuron. 74, 467–474.

7. Bartus, R. T., Dean, R. L3rd., Beer, B., Lippa, A. S., 1982. The cholinergic hypothesis of geriatric memory dysfunction. Science. 217, 408-414.

8. Beckman, M., Knox, K., Koneval, Z., Smith, C., Jayadev, S., Barker-Haliski, M., 2020. Loss of presenilin 2 age-dependently alters susceptibility to acute seizures and kindling acquisition. Neurobiol Dis. 136, 104719.

9. Bezzina, C., Verret, L., Juan, C., Remaud, J., Halley, H., Rampon, C., Dahan, L., 2015. Early onset of hypersynchronous network activity and expression of a marker of chronic seizures in the Tg2576 mouse model of Alzheimer’s disease. PLoS One. 10, e0119910.

10. Bliwise, D. L., Tinklenberg, J., Yesavage, J. A., Davies, H., Pursley, A. M., Petta, D. E., Widrow, L., Guilleminault, C., Zarcone, V. P., Dement, W. C., 1989. REM latency in Alzheimer’s disease. Biol Psychiatry. 25, 320–328.

11. Bragin, A., Jandó, G., Nádasdy, Z., van Landeghem, M., Buzsáki, G., 1995. Dentate EEG spikes and associated interneuronal population bursts in the hippocampal hilar region of the rat. J Neurophysiol. 73, 1691–1705.

12. Brown, R., Lam, A. D., Gonzalez-Sulser, A., Ying, A., Jones, M., Chou, R. C., Tzioras, M., Jordan, C. Y., Jedrasiak-Cape, I., Hemonnot, A. L., Abou Jaoude, M., Cole, A. J., Cash, S. S., Saito, T., Saido, T., Ribchester, R. R., Hashemi, K., Oren, I., 2018. Circadian and brain state modulation of network hyperexcitability in Alzheimer’s disease. eNeuro. 5.

13. Busche, M. A., Eichhoff, G., Adelsberger, H., Abramowski, D., Wiederhold, K. H., Haass, C., Staufenbiel, M., Konnerth, A., Garaschuk, O., 2008. Clusters of hyperactive neurons near amyloid plaques in a mouse model of Alzheimer’s disease. Science. 321, 1686–1689.

14. Buzsáki, G., 1986. Hippocampal sharp waves: Their origin and significance. Brain Res. 398, 242–252.

15. Buzsáki, G., Buhl, D. L., Harris, K. D., Csicsvari, J., Czéh, B., Morozov, A., 2003. Hippocampal network patterns of activity in the mouse. Neuroscience. 116, 201–211.

16. Cataldo, A. M., Petanceska, S., Peterhoff, C. M., Terio, N. B., Epstein, C. J., Villar, A., Carlson, E. J., Staufenbiel, M., Nixon, R. A., 2003. APP gene dosage modulates endosomal abnormalities of Alzheimer’s disease in a segmental trisomy 16 mouse model of Down syndrome. J Neurosci. 23, 6788–6792.

17. Choi, J. H., Berger, J. D., Mazzella, M. J., Morales-Corraliza, J., Cataldo, A. M., Nixon, R. A., Ginsberg, S. D., Levy, E., Mathews, P. M., 2009. Age-dependent dysregulation of brain amyloid precursor protein in the Ts65Dn down syndrome mouse model. J Neurochem. 110, 1818–1827.

18. Colom, L. V., 2006. Septal networks: Relevance to theta rhythm, epilepsy and Alzheimer’s disease. J Neurochem. 96, 609–623.

19. Corbett, B. F., You, J. C., Zhang, X., Pyfer, M. S., Tosi, U., Iascone, D. M., Petrof, I., Hazra, A., Fu, C. H., Stephens, G. S., Ashok, A. A., Aschmies, S., Zhao, L., Nestler, E. J., Chin, J., 2017. ΔFosB regulates gene expression and cognitive dysfunction in a mouse model of Alzheimer’s disease. Cell Rep. 20, 344–355.

20. Cortini, F., Cantoni, C., Villa, C., 2018. Epileptic seizures in autosomal dominant forms of Alzheimer’s disease. Seizure. 61, 4–7.

21. D’Rozario, A. L., Chapman, J. L., Phillips, C. L., Palmer, J. R., Hoyos, C. M., Mowszowski, L., Duffy, S. L., Marshall, N. S., Benca, R., Mander, B., Grunstein, R. R., Naismith, S. L., 2020. Objective measurement of sleep in mild cognitive impairment: a systematic review and meta-analysis. Sleep Medicine Reviews. 52, 101308.

22. DeKosky, S. T., Ikonomovic, M. D., Styren, S. D., Beckett, L., Wisniewski, S., Bennett, D. A., Cochran, E. J., Kordower, J. H., Mufson, E. J., 2002. Upregulation of choline acetyltransferase activity in hippocampus and frontal cortex of elderly subjects with mild cognitive impairment. Ann Neurol. 51, 145–155.

23. Desikan, S., Koser, D. E., Neitz, A., Monyer, H., 2018. Target selectivity of septal cholinergic neurons in the medial and lateral entorhinal cortex. Proc Natl Acad Sci U S A. 115, E2644–e2652.

24. Diamantaki, M., Frey, M., Berens, P., Preston-Ferrer, P., Burgalossi, A., 2016. Sparse activity of identified dentate granule cells during spatial exploration. eLife. 5, e20252.

25. Doppelbauer, A., Zeitlhofer, J., Zifko, U., Baumgartner, C., Mayr, N., Deecke, L., 1993. Occurrence of epileptiform activity in the routine EEG of epileptic patients. Acta Neurologica Scandinavica. 87, 345–352.

26. Duffy, A. M., Morales-Corraliza, J., Bermudez-Hernandez, K. M., Schaner, M. J., Magagna-Poveda, A., Mathews, P. M., Scharfman, H. E., 2015. Entorhinal cortical defects in Tg2576 mice are present as early as 2-4 months of age. Neurobiol Aging. 36, 134–148.

27. Dvorak, D., Chung, A., Park, E. H., Fenton, A. A., 2021. Dentate spikes and external control of hippocampal function. Cell Reports. 36, 109497.

28. Edwards, M., Robertson, N. P., 2018. Seizures in Alzheimer’s disease: is there more beneath the surface? J Neurol. 265, 226–228.

29. Ehrenberg, A. J., Kelberman, M. A., Liu, K. Y., Dahl, M. J., Weinshenker, D., Falgàs, N., Dutt, S., Mather, M., Ludwig, M., Betts, M. J., Winer, J. R., Teipel, S., Weigand, A. J., Eschenko, O., Hämmerer, D., Leiman, M., Counts, S. E., Shine, J. M., Robertson, I. H., Levey, A. I., Lancini, E., Son, G., Schneider, C., Egroo, M. V., Liguori, C., Wang, Q., Vazey, E. M., Rodriguez-Porcel, F., Haag, L., Bondi, M. W., Vanneste, S., Freeze, W. M., Yi, Y.-J., Maldinov, M., Gatchel, J., Satpati, A., Babiloni, C., Kremen, W. S., Howard, R., Jacobs, H. I. L., Grinberg, L. T., Priorities for research on neuromodulatory subcortical systems in Alzheimer’s disease: position paper from the NSS PIA of ISTAART. Alzheimer’s & Dementia.

30. Escorihuela, R. M., Vallina, I. F., Martínez-Cué, C., Baamonde, C., Dierssen, M., Tobeña, A., Flórez, J., Fernández-Teruel, A., 1998. Impaired short- and long-term memory in Ts65Dn mice, a model for down syndrome. Neurosci Lett. 247, 171–174.

31. Ezquerra, M., Lleó, A., Castellví, M., Queralt, R., Santacruz, P., Pastor, P., Molinuevo, J. L., Blesa, R., Oliva, R., 2003. A novel mutation in the PSEN2 gene (T430M) associated with variable expression in a family with early-onset Alzheimer disease. Arch Neurol. 60, 1149–1151.

32. Falangola, M. F., Nie, X., Ward, R., Dhiman, S., Voltin, J., Nietert, P. J., Jensen, J. H., 2021. Diffusion MRI detects basal forebrain cholinergic abnormalities in the 3xTG-AD mouse model of Alzheimer’s disease. J Magn Reson. 83, 1–13.

33. Faulkner, H. J., Arima, H., Mohamed, A., 2012. Latency to first interictal epileptiform discharge in epilepsy with outpatient ambulatory EEG. Clin Neurophysiol. 123, 1732–1735.

34. Fernández-Ruiz, A., Oliva, A., Soula, M., Rocha-Almeida, F., Nagy, G. A., Martin-Vazquez, G., Buzsáki, G., 2021. Gamma rhythm communication between entorhinal cortex and dentate gyrus neuronal assemblies. Science. 372.

35. Franklin, K. B. J., Paxinos, G., 1997. The mouse brain in stereotaxic coordinates. Academic Press, San Diego.

36. Friedman, A., Behrens, C. J., Heinemann, U., 2007. Cholinergic dysfunction in temporal lobe epilepsy. Epilepsia. 48 Suppl 5, 126–130.

37. Friedman, D., Honig, L. S., Scarmeas, N., 2012. Seizures and epilepsy in Alzheimer’s disease. CNS Neurosci Ther. 18, 285–294.

38. Galla, L., Redolfi, N., Pozzan, T., Pizzo, P., Greotti, E., 2020. Intracellular calcium dysregulation by the Alzheimer’s disease-linked protein presenilin 2. Int J Mol Sci. 21.

39. GoodSmith, D., Chen, X., Wang, C., Kim, S. H., Song, H., Burgalossi, A., Christian, K. M., Knierim, J. J., 2017. Spatial representations of granule cells and mossy cells of the dentate gyrus. Neuron. 93, 677–690.e675.

40. Gureviciene, I., Ishchenko, I., Ziyatdinova, S., Jin, N., Lipponen, A., Gurevicius, K., Tanila, H., 2019. Characterization of epileptic spiking associated with brain amyloidosis in APP/PS1 mice. Frontiers in Neurology. 10.

41. Guth, T. A., Kunz, L., Brandt, A., Dümpelmann, M., Klotz, K. A., Reinacher, P. C., Schulze-Bonhage, A., Jacobs, J., Schönberger, J., 2021. Interictal spikes with and without high-frequency oscillation have different single-neuron correlates. Brain. 144, 3078–3088.

42. Hamlett, E. D., Boger, H. A., Ledreux, A., Kelley, C. M., Mufson, E. J., Falangola, M. F., Guilfoyle, D. N., Nixon, R. A., Patterson, D., Duval, N., Granholm, A. C., 2016. Cognitive impairment, neuroimaging, and Alzheimer neuropathology in mouse models of Down syndrome. Curr Alzheimer Res. 13, 35–52.

43. Hampel, H., Mesulam, M. M., Cuello, A. C., Farlow, M. R., Giacobini, E., Grossberg, G. T., Khachaturian, A. S., Vergallo, A., Cavedo, E., Snyder, P. J., Khachaturian, Z. S., 2018. The cholinergic system in the pathophysiology and treatment of Alzheimer’s disease. Brain. 141, 1917–1933.

44. Hasselmo, M. E., Sarter, M., 2011. Modes and models of forebrain cholinergic neuromodulation of cognition. Neuropsychopharmacology. 36, 52–73.

45. Hauser, W. A., Morris, M. L., Heston, L. L., Anderson, V. E., 1986. Seizures and myoclonus in patients with Alzheimer’s disease. Neurology. 36, 1226–1230.

46. Head, E., Powell, D., Gold, B. T., Schmitt, F. A., 2012. Alzheimer’s disease in Down syndrome. Eur J Neurodegener Dis. 1, 353–364.

47. Headley, D. B., Kanta, V., Paré, D., 2017. Intra- and interregional cortical interactions related to sharp-wave ripples and dentate spikes. J Neurophysiol. 117, 556–565.

48. Hedreen, J. C., Struble, R. G., Whitehouse, P. J., Price, D. L., 1984. Topography of the magnocellular basal forebrain system in human brain. J Neuropathol Exp Neurol. 43, 1–21.

49. Herreman, A., Hartmann, D., Annaert, W., Saftig, P., Craessaerts, K., Serneels, L., Umans, L., Schrijvers, V., Checler, F., Vanderstichele, H., Baekelandt, V., Dressel, R., Cupers, P., Huylebroeck, D., Zwijsen, A., Van Leuven, F., De Strooper, B., 1999. Presenilin 2 deficiency causes a mild pulmonary phenotype and no changes in amyloid precursor protein processing but enhances the embryonic lethal phenotype of presenilin 1 deficiency. Proc Natl Acad Sci U S A. 96, 11872–11877.

50. Holtzman, D. M., Santucci, D., Kilbridge, J., Chua-Couzens, J., Fontana, D. J., Daniels, S. E., Johnson, R. M., Chen, K., Sun, Y., Carlson, E., Alleva, E., Epstein, C. J., Mobley, W. C., 1996. Developmental abnormalities and age-related neurodegeneration in a mouse model of Down syndrome. Proc Natl Acad Sci U S A. 93, 13333–13338.

51. Horak, P. C., Meisenhelter, S., Song, Y., Testorf, M. E., Kahana, M. J., Viles, W. D., Bujarski, K. A., Connolly, A. C., Robbins, A. A., Sperling, M. R., Sharan, A. D., Worrell, G. A., Miller, L. R., Gross, R. E., Davis, K. A., Roberts, D. W., Lega, B., Sheth, S. A., Zaghloul, K. A., Stein, J. M., Das, S. R., Rizzuto, D. S., Jobst, B. C., 2017. Interictal epileptiform discharges impair word recall in multiple brain areas. Epilepsia. 58, 373–380.

52. Horváth, A., Szűcs, A., Barcs, G., Kamondi, A., 2017. Sleep EEG detects epileptiform activity in Alzheimer’s disease with high sensitivity. J Alzheimers Dis. 56, 1175–1183.

53. Horvath, A. A., Papp, A., Zsuffa, J., Szucs, A., Luckl, J., Radai, F., Nagy, F., Hidasi, Z., Csukly, G., Barcs, G., Kamondi, A., 2021. Subclinical epileptiform activity accelerates the progression of Alzheimer’s disease: a long-term EEG study. Clin Neurophysiol. 132, 1982–1989.

54. Hsiao, K., Chapman, P., Nilsen, S., Eckman, C., Harigaya, Y., Younkin, S., Yang, F., Cole, G., 1996. Correlative memory deficits, Aβ elevation, and amyloid plaques in transgenic mice. Science. 274, 99–102.

55. Hyman, B. T., Kromer, L. J., Van Hoesen, G. W., 1987. Reinnervation of the hippocampal perforant pathway zone in Alzheimer’s disease. Ann Neurol. 21, 259–267.

56. Ishii, M., Wang, G., Racchumi, G., Dyke, J. P., Iadecola, C., 2014. Transgenic mice overexpressing amyloid precursor protein exhibit early metabolic deficits and a pathologically low leptin state associated with hypothalamic dysfunction in arcuate neuropeptide Y neurons. J Neurosci. 34, 9096–9106.

57. Jacobsen, J. S., Wu, C. C., Redwine, J. M., Comery, T. A., Arias, R., Bowlby, M., Martone, R., Morrison, J. H., Pangalos, M. N., Reinhart, P. H., Bloom, F. E., 2006. Early-onset behavioral and synaptic deficits in a mouse model of Alzheimer’s disease. Proc Natl Acad Sci U S A. 103, 5161–5166.

58. Jasper, H. H., Tessier, J., 1971. Acetylcholine liberation from cerebral cortex during paradoxical (REM) sleep. Science. 172, 601–602.

59. Jayadev, S., Case, A., Eastman, A. J., Nguyen, H., Pollak, J., Wiley, J. C., Möller, T., Morrison, R. S., Garden, G. A., 2010a. Presenilin 2 is the predominant γ-secretase in microglia and modulates cytokine release. PLoS One. 5, e15743.

60. Jayadev, S., Leverenz, J. B., Steinbart, E., Stahl, J., Klunk, W., Yu, C.-E., Bird, T. D., 2010b. Alzheimer’s disease phenotypes and genotypes associated with mutations in presenilin 2. Brain. 133, 1143–1154.

61. Jayadev, S., Leverenz, J. B., Steinbart, E., Stahl, J., Klunk, W., Yu, C. E., Bird, T. D., 2010c. Alzheimer’s disease phenotypes and genotypes associated with mutations in presenilin 2. Brain. 133, 1143–1154.

62. Jendryka, M., Palchaudhuri, M., Ursu, D., van der Veen, B., Liss, B., Kätzel, D., Nissen, W., Pekcec, A., 2019. Pharmacokinetic and pharmacodynamic actions of clozapine-N-oxide, clozapine, and compound 21 in DREADD-based chemogenetics in mice. Sci Rep. 9, 4522.

63. Jiang, N., Cupolillo, D., Grosjean, N., Muller, E., Deforges, S., Mulle, C., Amédée, T., 2021. Impaired plasticity of intrinsic excitability in the dentate gyrus alters spike transfer in a mouse model of Alzheimer’s disease. Neurobiol Dis. 154, 105345.

64. Jin, J., Cheng, J., Lee, K. W., Amreen, B., McCabe, K. A., Pitcher, C., Liebmann, T., Greengard, P., Flajolet, M., 2019. Cholinergic neurons of the medial septum are crucial for sensorimotor gating. J Neurosci. 39, 5234–5242.

65. Johnston, D., Brown, T. H., 1981. Giant synaptic potential hypothesis for epileptiform activity. Science. 211, 294–297.

66. Jung, M. W., McNaughton, B. L., 1993. Spatial selectivity of unit activity in the hippocampal granular layer. Hippocampus. 3, 165–182.

67. Kam, K., Duffy, A. M., Moretto, J., LaFrancois, J. J., Scharfman, H. E., 2016. Interictal spikes during sleep are an early defect in the Tg2576 mouse model of β-amyloid neuropathology. Sci Rep. 6, 20119.

68. Karoly, P. J., Freestone, D. R., Boston, R., Grayden, D. B., Himes, D., Leyde, K., Seneviratne, U., Berkovic, S., O’Brien, T., Cook, M. J., 2016. Interictal spikes and epileptic seizures: their relationship and underlying rhythmicity. Brain. 139, 1066–1078.

69. Kawarabayashi, T., Younkin, L. H., Saido, T. C., Shoji, M., Ashe, K. H., Younkin, S. G., 2001. Age-dependent changes in brain, CSF, and plasma amyloid-β protein in the Tg2576 transgenic mouse model of alzheimer’s disease. J Neurosci.l journal of the Society for Neuroscience. 21, 372-381.

70. Kent, B. A., Strittmatter, S. M., Nygaard, H. B., 2018. Sleep and EEG power spectral analysis in three transgenic mouse models of Alzheimer’s disease: APP/PS1, 3xTGAD, and Tg2576. J Alzheimers Dis. 64, 1325-1336.

71. Kim, K. R., Kim, Y., Jeong, H. J., Kang, J. S., Lee, S. H., Kim, Y., Lee, S. H., Ho, W. K., 2021. Impaired pattern separation in Tg2576 mice is associated with hyperexcitable dentate gyrus caused by Kv4.1 downregulation. Mol Brain. 14, 62.

72. Kim, S., Nam, Y., Jeong, Y. O., Park, H. H., Lee, S. K., Shin, S. J., Jung, H., Kim, B. H., Hong, S. B., Park, Y. H., Kim, J., Yu, J., Yoo, D. H., Park, S. H., Jeon, S. G., Moon, M., 2019. Topographical visualization of the reciprocal projection between the medial septum and the hippocampus in the 5xFAD mouse model of Alzheimer’s disease. Int J Mol Sci. 20.

73. Kipanyula, M. J., Contreras, L., Zampese, E., Lazzari, C., Wong, A. K., Pizzo, P., Fasolato, C., Pozzan, T., 2012. Ca^2+^ dysregulation in neurons from transgenic mice expressing mutant presenilin 2. Aging Cell. 11, 885–893.

74. Kleen, J. K., Scott, R. C., Holmes, G. L., Lenck-Santini, P. P., 2010. Hippocampal interictal spikes disrupt cognition in rats. Ann Neurol. 67, 250–257.

75. Klingner, M., Apelt, J., Kumar, A., Sorger, D., Sabri, O., Steinbach, J., Scheunemann, M., Schliebs, R., 2003. Alterations in cholinergic and non-cholinergic neurotransmitter receptor densities in transgenic Tg2576 mouse brain with β-amyloid plaque pathology. Int J Dev Neurosci. 21, 357–369.

76. Knox, K. M., Beckman, M., Smith, C. L., Jayadev, S., Barker-Haliski, M., 2023. Chronic seizures induce sex-specific cognitive deficits with loss of presenilin 2 function. Exp Neurol. 361, 114321.

77. Kokaia, M., Pratt, G. D., Elmér, E., Bengzon, J., Fritschy, J. M., Kokaia, Z., Lindvall, O., Mohler, H., 1994. Biphasic differential changes of GABAA receptor subunit mRNA levels in dentate gyrus granule cells following recurrent kindling-induced seizures. Brain Res Mol Brain Res. 23, 323–332.

78. Lahiri, D. K., Farlow, M. R., 2003. Review: Cholinesterase inhibitors have a modest effect on neuropsychiatric and functional outcomes in Alzheimer’s disease. Evid Based Ment Health. 6, 94.

79. Lam, A. D., Deck, G., Goldman, A., Eskandar, E. N., Noebels, J., Cole, A. J., 2017. Silent hippocampal seizures and spikes identified by foramen ovale electrodes in Alzheimer’s disease. Nat Med. 23, 678–680.

80. Lam, A. D., Noebels, J., 2020. Night watch on the titanic: detecting early signs of epileptogenesis in Alzheimer disease. Epilepsy Curr. 20, 369–374.

81. Lam, A. D., Sarkis, R. A., Pellerin, K. R., Jing, J., Dworetzky, B. A., Hoch, D. B., Jacobs, C. S., Lee, J. W., Weisholtz, D. S., Zepeda, R., Westover, M. B., Cole, A. J., Cash, S. S., 2020. Association of epileptiform abnormalities and seizures in Alzheimer disease. Neurology. 95, e2259–e2270.

82. Lehmann, L., Lo, A., Knox, K. M., Barker-Haliski, M., 2021. Alzheimer’s disease and epilepsy: a perspective on the opportunities for overlapping therapeutic innovation. Neurochem Res. 46, 1895–1912.

83. Liedorp, M., Stam, C. J., van der Flier, W. M., Pijnenburg, Y. A., Scheltens, P., 2010. Prevalence and clinical significance of epileptiform EEG discharges in a large memory clinic cohort. Dement Geriatr Cogn Disord. 29, 432–437.

84. Lisgaras, C. P., Oliva, A., McKenzie, S., LaFrancois, J., Siegelbaum, S. A., Scharman, H. E., 2023. Hippocampal area CA2 controls seizure dynamics, interictal EEG abnormalities and social comorbidity in mouse models of temporal lobe epilepsy. BioRxiv.

85. Lisgaras, C. P., Scharfman, H. E., 2022a. High frequency oscillations (250-500 Hz) in animal models of Alzheimer’s disease and two animal models of epilepsy. Epilepsia. 231–246

86. Lisgaras, C. P., Scharfman, H. E., 2022b. Robust chronic convulsive seizures, high frequency oscillations, and human seizure onset patterns in an intrahippocampal kainic acid model in mice. Neurobiol Dis. 166:105637.

87. Lleó, A., Blesa, R., Gendre, J., Castellví, M., Pastor, P., Queralt, R., Oliva, R., 2001. A novel presenilin 2 gene mutation (D439A) in a patient with early-onset Alzheimer’s disease. Neurology. 57, 1926–1928.

88. Manvich, D. F., Webster, K. A., Foster, S. L., Farrell, M. S., Ritchie, J. C., Porter, J. H., Weinshenker, D., 2018. The DREADD agonist clozapine n-oxide (CNO) is reverse-metabolized to clozapine and produces clozapine-like interoceptive stimulus effects in rats and mice. Sci Rep. 8, 3840.

89. Maslarova, A., Salar, S., Lapilover, E., Friedman, A., Veh, R. W., Heinemann, U., 2013. Increased susceptibility to acetylcholine in the entorhinal cortex of pilocarpine-treated rats involves alterations in KCNQ channels. Neurobiol Dis. 56, 14–24.

90. Matsumoto, H., Marsan, C. A., 1964. Cortical cellular phenomena in experimental epilepsy: Interictal manifestations. Experimental Neurology. 9, 286–304.

91. Mikroulis, A., Lisgaras, C. P., Psarropoulou, C., 2018. Immature status epilepticus: *In vitro* models reveal differences in cholinergic control and HFO properties of adult CA3 interictal discharges in temporal vs septal hippocampus. Neuroscience. 369, 386–398.

92. Minkeviciene, R., Rheims, S., Dobszay, M. B., Zilberter, M., Hartikainen, J., Fülöp, L., Penke, B., Zilberter, Y., Harkany, T., Pitkänen, A., Tanila, H., 2009. Amyloid β-induced neuronal hyperexcitability triggers progressive epilepsy. J Neurosci. 29, 3453–3462.

93. Mufson, E. J., Counts, S. E., Perez, S. E., Ginsberg, S. D., 2008. Cholinergic system during the progression of Alzheimer’s disease: Therapeutic implications. Expert Rev Neurother. 8, 1703–1718.

94. Muldoon, S. F., Villette, V., Tressard, T., Malvache, A., Reichinnek, S., Bartolomei, F., Cossart, R., 2015. GABAergic inhibition shapes interictal dynamics in awake epileptic mice. Brain. 138, 2875–2890.

95. Navarrete, M., Alvarado-Rojas, C., Le Van Quyen, M., Valderrama, M., 2016. Ripplelab: A comprehensive application for the detection, analysis and classification of high frequency oscillations in electroencephalographic signals. PLoS One. 11, e0158276.

96. Ohm, T. G., The dentate gyrus in Alzheimer’s disease. In: H. E. Scharfman, (Ed.), Progress in brain research. New York: Elsevier; 2007. p. 723–740.

97. Ohno, M., Sametsky, E. A., Younkin, L. H., Oakley, H., Younkin, S. G., Citron, M., Vassar, R., Disterhoft, J. F., 2004. BACE1 deficiency rescues memory deficits and cholinergic dysfunction in a mouse model of Alzheimer’s disease. Neuron. 41, 27–33.

98. Oliva, A., Fernandez-Ruiz, A., Buzsaki, G., Berenyi, A., 2016. Role of hippocampal CA2 region in triggering sharp-wave ripples. Neuron. 91, 1342–1355.

99. Pace-Schott, E. F., Hobson, J. A., 2002. The neurobiology of sleep: genetics, cellular physiology and subcortical networks. Nat Rev Neurosci. 3, 591–605.

100. Palop, J. J., Chin, J., Roberson, E. D., Wang, J., Thwin, M. T., Bien-Ly, N., Yoo, J., Ho, K. O., Yu, G. Q., Kreitzer, A., Finkbeiner, S., Noebels, J. L., Mucke, L., 2007. Aberrant excitatory neuronal activity and compensatory remodeling of inhibitory hippocampal circuits in mouse models of Alzheimer’s disease. Neuron. 55, 697–711.

101. Palop, J. J., Jones, B., Kekonius, L., Chin, J., Yu, G. Q., Raber, J., Masliah, E., Mucke, L., 2003. Neuronal depletion of calcium-dependent proteins in the dentate gyrus is tightly linked to Alzheimer’s disease-related cognitive deficits. Proc Natl Acad Sci U S A. 100, 9572–9577.

102. Palop, J. J., Mucke, L., 2009. Epilepsy and cognitive impairments in Alzheimer disease. Arch Neurol. 66, 435–440.

103. Petersen, P. C., Siegle, J. H., Steinmetz, N. A., Mahallati, S., Buzsáki, G., 2021. Cellexplorer: a framework for visualizing and characterizing single neurons. Neuron. 109, 3594–3608.e3592.

104. Pofahl, M., Nikbakht, N., Haubrich, A. N., Nguyen, T., Masala, N., Distler, F., Braganza, O., Macke, J. H., Ewell, L. A., Golcuk, K., Beck, H., 2021. Synchronous activity patterns in the dentate gyrus during immobility. Elife. 10.

105. Reinholdt, L. G., Ding, Y., Gilbert, G. J., Czechanski, A., Solzak, J. P., Roper, R. J., Johnson, M. T., Donahue, L. R., Lutz, C., Davisson, M. T., 2011. Molecular characterization of the translocation breakpoints in the Down syndrome mouse model Ts65Dn. Mamm Genome. 22, 685–691.

106. Rossi, J., Balthasar, N., Olson, D., Scott, M., Berglund, E., Lee, C. E., Choi, M. J., Lauzon, D., Lowell, B. B., Elmquist, J. K., 2011. Melanocortin-4 receptors expressed by cholinergic neurons regulate energy balance and glucose homeostasis. Cell Metab. 13, 195–204.

107. Roth, B. L., 2016. DREADDs for neuroscientists. Neuron. 89, 683–694.

108. Rozalem Aranha, M., Iulita, M. F., Montal, V., Pegueroles, J., Bejanin, A., Vaqué-Alcázar, L., Grothe, M. J., Carmona-Iragui, M., Videla, L., Benejam, B., Arranz, J., Padilla, C., Valldeneu, S., Barroeta, I., Altuna, M., Fernández, S., Ribas, L., Valle-Tamayo, N., Alcolea, D., González-Ortiz, S., Bargalló, N., Zetterberg, H., Blennow, K., Blesa, R., Wisniewski, T., Busciglio, J., Cuello, A. C., Lleó, A., Fortea, J., 2023. Basal forebrain atrophy along the Alzheimer’s disease continuum in adults with Down syndrome. Alzheimers Dement.

109. Salehi, A., Ashford, J. W., Mufson, E. J., 2016. The link between Alzheimer’s disease and Down syndrome. A historical perspective. Curr Alzheimer Res. 13, 2–6.

110. Salehi, A., Delcroix, J. D., Belichenko, P. V., Zhan, K., Wu, C., Valletta, J. S., Takimoto-Kimura, R., Kleschevnikov, A. M., Sambamurti, K., Chung, P. P., Xia, W., Villar, A., Campbell, W. A., Kulnane, L. S., Nixon, R. A., Lamb, B. T., Epstein, C. J., Stokin, G. B., Goldstein, L. S., Mobley, W. C., 2006. Increased APP expression in a mouse model of Down’s syndrome disrupts NGF transport and causes cholinergic neuron degeneration. Neuron. 51, 29–42.

111. Sammaritano, M., Gigli, G. L., Gotman, J., 1991. Interictal spiking during wakefulness and sleep and the localization of foci in temporal lobe epilepsy. Neurology. 41, 290–297.

112. Sanchez, P. E., Zhu, L., Verret, L., Vossel, K. A., Orr, A. G., Cirrito, J. R., Devidze, N., Ho, K., Yu, G. Q., Palop, J. J., Mucke, L., 2012. Levetiracetam suppresses neuronal network dysfunction and reverses synaptic and cognitive deficits in an Alzheimer’s disease model. Proc Natl Acad Sci U S A. 109, E2895–2903.

113. Saper, C. B., Chou, T. C., Scammell, T. E., 2001. The sleep switch: hypothalamic control of sleep and wakefulness. Trends Neurosci. 24, 726–731.

114. Scarmeas, N., Honig, L. S., Choi, H., Cantero, J., Brandt, J., Blacker, D., Albert, M., Amatniek, J. C., Marder, K., Bell, K., Hauser, W. A., Stern, Y., 2009. Seizures in Alzheimer disease: who, when, and how common? Arch Neurol. 66, 992–997.

115. Scharfman, H. E., 2012. “Untangling” Alzheimer’s disease and epilepsy. Epilepsy Curr. 12, 178–183.

116. Schwartzkroin, P. A., Prince, D. A., 1977. Penicillin-induced epileptiform activity in the hippocampal *in vitro* prepatation. Ann Neurol. 1, 463–469.

117. Senzai, Y., Buzsáki, G., 2017. Physiological properties and behavioral correlates of hippocampal granule cells and mossy cells. Neuron. 93, 691–704.e695.

118. Smith, K. S., Bucci, D. J., Luikart, B. W., Mahler, S. V., 2016. DREADDs: use and application in behavioral neuroscience. Behav Neurosci. 130, 137–155.

119. Sperling, R. A., Jack, C. R., Jr., Aisen, P. S., 2011. Testing the right target and right drug at the right stage. Sci Transl Med. 3, 111cm133.

120. Staley, K. J., Dudek, F. E., 2006. Interictal spikes and epileptogenesis. Epilepsy Currents. 6, 199–202.

121. Stickgold, R., 2005. Sleep-dependent memory consolidation. Nature. 437, 1272–1278.

122. Taheri, S., Zeitzer, J. M., Mignot, E., 2002. The role of hypocretins (orexins) in sleep regulation and narcolepsy. Annu Rev Neurosci. 25, 283–313.

123. Trinh, N. H., Hoblyn, J., Mohanty, S., Yaffe, K., 2003. Efficacy of cholinesterase inhibitors in the treatment of neuropsychiatric symptoms and functional impairment in Alzheimer disease: A meta-analysis. JAMA. 289, 210–216.

124. Ulbert, I., Heit, G., Madsen, J., Karmos, G., Halgren, E., 2004. Laminar analysis of human neocortical interictal spike generation and propagation: current source density and multiunit analysis *in vivo*. Epilepsia. 45 Suppl 4, 48–56.

125. Van Dort, C. J., Zachs, D. P., Kenny, J. D., Zheng, S., Goldblum, R. R., Gelwan, N. A., Ramos, D. M., Nolan, M. A., Wang, K., Weng, F. J., Lin, Y., Wilson, M. A., Brown, E. N., 2015. Optogenetic activation of cholinergic neurons in the PPT or LDT induces REM sleep. Proc Natl Acad Sci U S A. 112, 584–589.

126. Vandecasteele, M., Varga, V., Berényi, A., Papp, E., Barthó, P., Venance, L., Freund, T. F., Buzsáki, G., 2014. Optogenetic activation of septal cholinergic neurons suppresses sharp wave ripples and enhances theta oscillations in the hippocampus. Proc Natl Acad Sci U S A. 111, 13535–13540.

127. Vazquez, J., Baghdoyan, H. A., 2001. Basal forebrain acetylcholine release during REM sleep is significantly greater than during waking. Am J of Physiol Regul Integr Comp Physiol. 280, R598–R601.

128. Vitiello, M. V., Bokan, J. A., Kukull, W. A., Muniz, R. L., Smallwood, R. G., Prinz, P. N., 1984. Rapid eye movement sleep measures of Alzheimer’s-type dementia patients and optimally healthy aged individuals. Biol Psychiatry. 19, 721–734.

129. Vossel, K., Ranasinghe, K. G., Beagle, A. J., La, A., Ah Pook, K., Castro, M., Mizuiri, D., Honma, S. M., Venkateswaran, N., Koestler, M., Zhang, W., Mucke, L., Howell, M. J., Possin, K. L., Kramer, J. H., Boxer, A. L., Miller, B. L., Nagarajan, S. S., Kirsch, H. E., 2021. Effect of levetiracetam on cognition in patients with Alzheimer disease with and without epileptiform activity: a randomized clinical trial. JAMA Neurol. 78, 1345–1354.

130. Vossel, K. A., Beagle, A. J., Rabinovici, G. D., Shu, H., Lee, S. E., Naasan, G., Hegde, M., Cornes, S. B., Henry, M. L., Nelson, A. B., Seeley, W. W., Geschwind, M. D., Gorno-Tempini, M. L., Shih, T., Kirsch, H. E., Garcia, P. A., Miller, B. L., Mucke, L., 2013. Seizures and epileptiform activity in the early stages of Alzheimer disease. JAMA Neurol. 70, 1158–1166.

131. Vossel, K. A., Ranasinghe, K. G., Beagle, A. J., Mizuiri, D., Honma, S. M., Dowling, A. F., Darwish, S. M., Van Berlo, V., Barnes, D. E., Mantle, M., Karydas, A. M., Coppola, G., Roberson, E. D., Miller, B. L., Garcia, P. A., Kirsch, H. E., Mucke, L., Nagarajan, S. S., 2016. Incidence and impact of subclinical epileptiform activity in Alzheimer’s disease. Ann Neurol. 80, 858–870.

132. Vossel, K. A., Tartaglia, M. C., Nygaard, H. B., Zeman, A. Z., Miller, B. L., 2017. Epileptic activity in Alzheimer’s disease: causes and clinical relevance. Lancet Neurol. 16, 311–322.

133. Whissell, P. D., Tohyama, S., Martin, L. J., 2016. The use of DREADDs to deconstruct behavior. Front Genet. 7, 70.

134. Wisniewski, K. E., Wisniewski, H. M., Wen, G. Y., 1985. Occurrence of neuropathological changes and dementia of Alzheimer’s disease in Down’s syndrome. Ann Neurol. 17, 278–282.

135. Wisor, J. P., Edgar, D. M., Yesavage, J., Ryan, H. S., McCormick, C. M., Lapustea, N., Murphy, G. M., Jr., 2005. Sleep and circadian abnormalities in a transgenic mouse model of Alzheimer’s disease: a role for cholinergic transmission. Neuroscience. 131, 375–385.

136. Zarea, A., Charbonnier, C., Rovelet-Lecrux, A., Nicolas, G., Rousseau, S., Borden, A., Pariente, J., Le Ber, I., Pasquier, F., Formaglio, M., Martinaud, O., Rollin-Sillaire, A., Sarazin, M., Croisile, B., Boutoleau-Bretonnière, C., Ceccaldi, M., Gabelle, A., Chamard, L., Blanc, F., Sellal, F., Paquet, C., Campion, D., Hannequin, D., Wallon, D., 2016. Seizures in dominantly inherited Alzheimer disease. Neurology. 87, 912–919.

137. Zatti, G., Burgo, A., Giacomello, M., Barbiero, L., Ghidoni, R., Sinigaglia, G., Florean, C., Bagnoli, S., Binetti, G., Sorbi, S., Pizzo, P., Fasolato, C., 2006. Presenilin mutations linked to familial Alzheimer’s disease reduce endoplasmic reticulum and golgi apparatus calcium levels. Cell Calcium. 39, 539–550.

138. Zatti, G., Ghidoni, R., Barbiero, L., Binetti, G., Pozzan, T., Fasolato, C., Pizzo, P., 2004. The presenilin 2 M239I mutation associated with familial Alzheimer’s disease reduces Ca^2+^ release from intracellular stores. Neurobiol Dis. 15, 269–278.

139. Zhang, Y., Cao, L., Varga, V., Jing, M., Karadas, M., Li, Y., Buzsáki, G., 2021. Cholinergic suppression of hippocampal sharp-wave ripples impairs working memory. Proceedings of the National Academy of Sciences. 118, e2016432118.

140. Zhang, Y., Ren, R., Yang, L., Zhang, H., Shi, Y., Okhravi, H. R., Vitiello, M. V., Sanford, L. D., Tang, X., 2022. Sleep in Alzheimer’s disease: a systematic review and meta-analysis of polysomnographic findings. Translational Psychiatry. 12, 136.

141. Zhu, H., Aryal, D. K., Olsen, R. H., Urban, D. J., Swearingen, A., Forbes, S., Roth, B. L., Hochgeschwender, U., 2016. Cre-dependent DREADD (designer receptors exclusively activated by designer drugs) mice. Genesis. 54, 439–446.

